# Concerted SUMO-targeted ubiquitin ligase activities of TOPORS and RNF4 are essential for stress management and cell proliferation

**DOI:** 10.1101/2023.12.20.572718

**Authors:** Julio C.Y. Liu, Leena Ackermann, Saskia Hoffmann, Zita Gál, Ivo A. Hendriks, Charu Jain, Louise Morlot, Michael H. Tatham, Gian-Luca McLelland, Ronald T. Hay, Michael Lund Nielsen, Thijn Brummelkamp, Peter Haahr, Niels Mailand

## Abstract

Protein SUMOylation provides a principal driving force for cellular stress responses including DNA-protein crosslink (DPC) repair and arsenic-induced PML body degradation. In genome-scale screens, we identified the human E3 ligase TOPORS as a key effector of SUMO-dependent DPC resolution. We demonstrate that TOPORS promotes DPC repair by functioning as a SUMO-targeted ubiquitin ligase (STUbL) for DPCs, combining ubiquitin ligase activity through its RING domain with poly-SUMO chain binding via a cluster of SUMO-interacting motifs, analogous to the STUbL RNF4. Surprisingly, the STUbL activities of TOPORS and RNF4 are both required for SUMO-dependent DPC repair, PML degradation and other stress responses, making overlapping and distinct contributions to ubiquitin chain formation on SUMOylated targets to enable p97/VCP unfoldase recruitment. Combined loss of TOPORS and RNF4 is synthetic lethal even in unstressed cells, leading to defective clearance of SUMOylated proteins from chromatin accompanied by cell cycle arrest and apoptosis. Together, our findings establish TOPORS as a novel STUbL whose concerted action with RNF4 defines a general mechanistic principle in crucial cellular processes governed by direct SUMO-ubiquitin crosstalk.

**Highlights:**

- The RING E3 ligase TOPORS is required for SUMO-dependent DPC repair
- TOPORS is a novel SUMO-targeted ubiquitin ligase (STUbL)
- TOPORS promotes multiple STUbL-driven processes in conjunction with RNF4
- Combined TOPORS and RNF4 loss is synthetic lethal in human cells

## Introduction

Protein modification by members of the family of small ubiquitin-like modifier (SUMO) polypeptides impacts thousands of cellular targets and constitutes an important regulatory signaling mechanism in numerous aspects of biology, especially cellular stress responses ^1,2^. Protein SUMOylation proceeds via a three-step enzymatic cascade involving E1, E2 and E3 enzymes, similar to the machinery underlying protein ubiquitylation. However, whereas the ubiquitin signaling network displays a high degree of complexity, comprising hundreds of components, the SUMO modification machinery is comparatively simple, involving single E1 and E2 enzymes, and less than a dozen known or reported SUMO E3 ligases ^1^. While SUMOylation *per se* is a non-proteolytic modification, extensive crosstalk between SUMO and ubiquitin signals occurs in cells and positions SUMOylation to effectively serve as a protein modification that instructs the proteasomal degradation of many SUMO targets, brought about by their SUMO-dependent ubiquitylation. To a large extent, this direct coupling between SUMO and ubiquitin is mediated by SUMO-targeted ubiquitin ligases (STUbLs), which combine E3 ubiquitin ligase activity with SUMO-interacting motifs (SIMs), thereby enabling them to specifically recognize and ubiquitylate SUMOylated proteins ^1,3^. STUbLs typically contain multiple SIMs, imparting selectivity for proteins modified by poly-SUMOylation, which predominantly involves SUMO2/3 that unlike SUMO1 undergoes efficient chain formation, similar to ubiquitin ^1^. Reminiscent of STUbLs, the deubiquitinase USP7 has been reported to function as a SUMO-targeted ubiquitin protease counteracting SUMO-dependent protein ubiquitylation and degradation ^4^.

Two STUbLs, RNF4 and RNF111, which differ with respect to their preference for recognizing different SUMO chain configurations, are currently known in mammalian cells ^5–9^. These proteins have key roles in genome stability maintenance processes, mediated in part by their functions in regulating protein interactions with chromatin by promoting extraction of factors that undergo SUMOylation upon their association with chromatin ^1,3,10^. An extreme case of this regulatory principle is seen for DNA-protein crosslinks (DPCs), in which the adducted protein is covalently trapped on DNA and consequently becomes extensively SUMOylated ^11^. DPCs are bulky and highly toxic DNA lesions produced by many chemotherapeutic agents that undermine genome integrity by obstructing essential DNA-associated processes ^12,13^. Previous work by us and others showed that outside the context of DNA replication, resolution of DPCs critically relies on SUMOylation of the protein adducts, which triggers their subsequent ubiquitylation by RNF4 to promote proteolytic processing of the trapped proteins via the p97/VCP unfoldase and the proteasome ^11,14–16^. This mode of regulation is prominently observed for DPCs involving the maintenance DNA methyltransferase DNMT1, which can be efficiently generated in a post-replicative manner by the cytosine analog 5-Aza-2’-deoxycytidine (5-AzadC), which following its incorporation into genomic DNA during chromosome replication forms a DPC with the DNMT1 catalytic cysteine upon DNA methylation re-establishment ^11,14,17^. Importantly, 5-AzadC-induced DNMT1 adducts not only provide a defined model for studying mechanisms of SUMO-dependent resolution of DPCs, as 5-AzadC (also known as Decitabine) and related DNA methyltransferase inhibitors are used in the clinic for treatment of patients with acute myeloid leukemia and myelodysplastic syndrome ^18^. Recent findings showed that combining 5-AzadC and SUMO inhibitor treatment synergizes in reducing lymphoma tumor cell growth *in vitro* and *in vivo* ^19^, consistent with the requirement for SUMOylation in promoting resolution of 5-AzadC-induced DPCs. Direct SUMO-ubiquitin crosstalk also has a well-established critical role in promoting proteasomal degradation of PML bodies in response to arsenic trioxide treatment, which induces SUMOylation of the PML protein and thereby renders it susceptible to the STUbL activity of RNF4 ^8,9^. This has important ramifications for the treatment of acute promyelocytic leukemias caused by a chromosomal translocation fusing the *PML* and retinoic acid receptor alpha (*RARA*) genes, most patients of which can now be cured by combination therapy involving arsenic and retinoic acid ^20^. From these and other studies, it is becoming clear that STUbLs are crucial effectors of a growing range of SUMO-regulated cellular stress responses, whose pharmacological intervention have a demonstrated potential in cancer therapeutics. However, whether mammalian genomes encode STUbLs in addition to RNF4 and RNF111 is unclear.

In this study, we discovered that the E3 ligase protein TOPORS functions as a novel STUbL in human cells by virtue of ubiquitin ligase activity and a poly-SUMO-binding SIM cluster. We demonstrate that the STUbL activities of TOPORS and RNF4 act in conjunction to promote DPC resolution, PML body degradation and other SUMO-driven stress responses by promoting efficient multi-linkage poly-ubiquitylation of SUMOylated targets to facilitate their processing via the p97-proteasome pathway. Moreover, we reveal a strong synthetic lethal interaction between RNF4 and TOPORS or the deubiquitinase USP7, whose activity is critically required for sustaining TOPORS stability, characterized by accumulation of hyper-SUMOylated proteins on chromatin accompanied by defective cell cycle progression and apoptosis. Collectively, our findings reveal an unexpected mechanistic complexity of STUbL-driven processes involving concerted TOPORS and RNF4 E3 ligase activities as a general underlying principle in both fundamental cellular processes and stress responses governed by SUMO-ubiquitin crosstalk.

## Results

### Genome-wide screens reveal an essential role of TOPORS in SUMO-dependent DPC repair

We previously demonstrated that the total cellular pool of DNMT1 is rapidly and quantitatively depleted following 5-AzadC treatment due to covalent trapping on DNA and subsequent SUMO- and RNF4-mediated proteasomal turnover ^11,14^. Taking advantage of this notion, we carried out two complementary genome-wide insertional gene-trap mutagenesis screens with ultra-deep resolution in human HAP1 cells to identify regulators of SUMO-dependent DNMT1 DPC resolution.

Mutagenized cells subjected to 5-AzadC or mock treatment were fixed and stained with a specific DNMT1 antibody, and cell populations with relatively high or low antibody staining were isolated by FACS and analyzed by deep sequencing (**Figure 1A; Figure S1A**). The screens revealed a range of genes that were selectively enriched for disruptive gene-trap mutations in cells with high DNMT1 signal following 5-AzadC treatment, but not in mock-treated cells, indicative of a potential DNMT1 DPC repair defect (**Figure 1B; Figure S1B; Table S1**). Among these, both *RNF4* and *SUMO2* scored as specific negative regulators of DNMT1 abundance in 5-AzadC-treated cells, consistent with their key roles in DNMT1 DPC degradation ^14^, whereas the antibody target encoded by *DNMT1* was the strongest positive regulator as expected (**Figure 1B**). The *DCK* gene encoding deoxycytidine kinase (DCK) that is required for 5-AzadC incorporation into genomic DNA (and thus DNMT1 DPC formation) by converting it to 5-AzadCMP ^21^, also scored as a prominent negative regulator (**Figure 1B**), further supporting the validity of our genetic screen. Interestingly, we noted that insertions in the gene encoding TOPORS, a RING-type E3 ligase protein, were strongly enriched among cells showing defective DNMT1 degradation upon 5-AzadC treatment (**Figure 1B**). To confirm the involvement of TOPORS in proteolytic DNMT1 DPC repair, we used previously established chromatin fractionation- and quantitative imaging-based assays to monitor the impact of TOPORS knockout (KO) (**Figure S1C**) or depletion by independent siRNAs (**Figure S1D**) on DNMT1 DPC resolution kinetics following 5-AzadC treatment ^14^. Consistent with the screen results, TOPORS KO in independent HAP1 clones strongly impaired 5-AzadC-induced DNMT1 DPC degradation (**Figure 1C**). Likewise, knockdown of TOPORS in U2OS osteosarcoma cells led to a severe defect in DNMT1 DPC removal (**Figure 1D,E; Figure S1E**). TOPORS deficiency also impaired the timely clearance of SUMO2/3 foci induced by treatment with formaldehyde, another potent inducer of DPCs that are processed via SUMOylation ^11^ (**Figure S1F,G**). Supporting a direct involvement of TOPORS in processing SUMOylated DPCs, we found that TOPORS interacted with DNMT1 in a 5-AzadC- and SUMO-dependent manner, mirroring the behavior of RNF4 (**Figure 1F**). Furthermore, consistent with a role for TOPORS in promoting turnover of 5-AzadC- and formaldehyde-induced DPCs, cells lacking TOPORS displayed hypersensitivity to treatment with these agents (**Figure 1G; Figure S1H**), similar to our previous findings for RNF4 ^14^. Additional components of the ubiquitin network including *USP7* and *UBE2K* were also enriched as significant negative regulators of DNMT1 abundance upon 5-AzadC treatment (**Figure 1B; Table S1**). Interestingly, recent proteomic studies suggested acute and highly selective loss of TOPORS expression upon treatment of cells with inhibitors of the deubiquitinase USP7 ^22,23^, and we confirmed that a specific USP7 inhibitor, FT671 ^24^ (referred to as USP7i), caused a rapid decline in TOPORS abundance whereas RNF4 levels were not affected (**Figure 1H**). Consistently, USP7i delayed DNMT1 DPC turnover and hypersensitized cells to 5-AzadC (**Figure 1I,J; Figure S1I**), similar to the effect of TOPORS depletion. However, unlike TOPORS and RNF4, USP7 did not display 5-AzadC- and SUMO-dependent interaction with DNMT1 (**Figure 1F**), suggesting that it may function indirectly in SUMO-dependent DPC resolution by underpinning TOPORS stability rather than acting directly at DPCs. Indeed, TOPORS, but not RNF4, interacted with USP7 and was modified by auto-ubiquitylation that was efficiently antagonized by recombinant USP7 (**Figure S1J-L**). Collectively, these data identify TOPORS as a critical novel factor in SUMO-dependent DPC resolution and suggest that USP7 is required for this response by sustaining TOPORS expression via its deubiquitinase activity.

**Figure 1.**
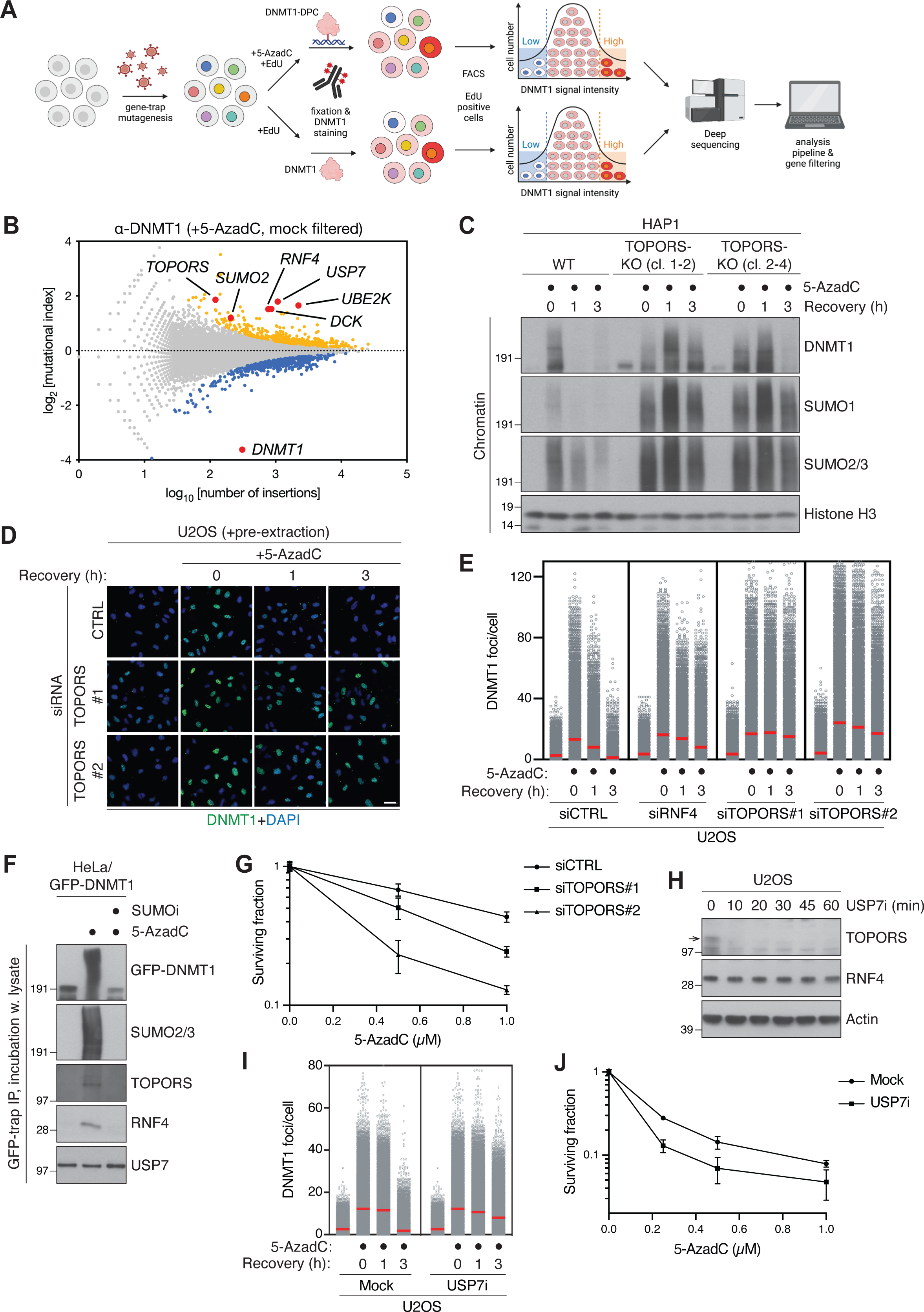
Genome-wide screens reveal an essential role of TOPORS in SUMO-dependent DPC repair. **A.** Workflow schematic of FACS-based haploid genetic screens for DNMT1 abundance. **B.** Screen for DNMT1 abundance in EdU-positive cells following co-treatment with 5-AzadC and EdU for 2 h, presented as a fishtail plot in which genes are plotted according to their mutational index (y-axis) and the total number of gene-trap insertions in the gene (x-axis). Positive and negative regulators of DNMT1 abundance are labeled blue and yellow, respectively (two-sided Fisher’s exact test, false discovery rate (FDR) corrected p≤0.05; non-significant genes are shown in grey). Genes that also score as significant negative or positive DNMT1 regulators in a mock screen (**Figure S1B**) were filtered out, except for *DNMT1*. **C.** HAP1 WT or TOPORS-KO cell lines released from single-round thymidine synchronization in early S phase were treated with 5-AzadC for 30 min, then washed and collected at the indicated times. Chromatin-enriched fractions were immunoblotted with indicated antibodies. **D.** Representative images of U2OS cells transfected with indicated siRNAs, treated as in (C) and pre-extracted and immunostained with DNMT1 antibody. **E.** DNMT1 foci formation in cells in (D) was analyzed by quantitative image-based cytometry (QIBC) (red bars, mean; >8,300 cells analyzed per condition). Data are representative of 3 independent experiments. **F.** HeLa cells stably expressing GFP-DNMT1 were synchronized in early S phase by double thymidine block and release, treated with 5-AzadC and/or SUMOi for 30 min, collected and subjected to GFP immunoprecipitation (IP) under denaturing conditions. Following stringent washing, immobilized GFP-DNMT1 complexes were incubated with whole cell lysate from parental HeLa cells, washed extensively and immunoblotted with indicated antibodies. **G.** Clonogenic survival of 5-AzadC-treated U2OS cells transfected with indicated siRNAs (mean±SEM; *n*=3 independent experiments). **H.** Immunoblot analysis of U2OS cells treated with USP7i for the indicated times. Arrow indicates the band corresponding to endogenous TOPORS. **I.** U2OS cells released from single thymidine synchronization in early S phase were treated with SUMOi and 5-AzadC for 30 min, then washed, released into medium containing USP7i or not, and collected at the indicated times. Cells were immunostained with DNMT1 antibody and analyzed by QIBC as in (D) (red bars, mean; >20,000 cells analyzed per condition). Data are representative of 3 independent experiments. **J.** Clonogenic survival of 5-AzadC-treated U2OS cells in the presence or absence of USP7i (mean±SEM; *n*=3 independent experiments).

### TOPORS functions as a SUMO-targeted ubiquitin ligase in DPC repair

We next set out to address how TOPORS promotes SUMO-dependent DPC resolution. While TOPORS contains a conserved N-terminal RING domain (**Figure 2A**) its precise enzymatic function remains unclear as, unusually, TOPORS has been reported to function as an E3 ligase for both ubiquitin and SUMO ^25–29^. We therefore asked how TOPORS impacts DNMT1 DPC SUMOylation and ubiquitylation levels, using stringent nanobody-based isolation of endogenous DNMT1 under denaturing conditions. Interestingly, TOPORS knockdown led to a marked decrease in DNMT1 DPC ubiquitylation whereas the level of DNMT1 DPC SUMOylation was greatly enhanced (**Figure 2B; Figure S2A**), reminiscent of our previous findings for RNF4 ^14^. Co-depletion of RNF4 and TOPORS, the effect of which could only be assessed at early time points after siRNA transfection due to synthetic lethality as detailed below (see Figure 5), reduced DNMT1 DPC ubiquitylation but not SUMOylation to background levels despite individual depletion of either protein for a short, 40-h duration only modestly decreased DPC ubiquitylation (**Figure 2C**) unlike the impact of TOPORS knockdown for a longer (72-h) period (**Figure 2B**).

**Figure 2.**
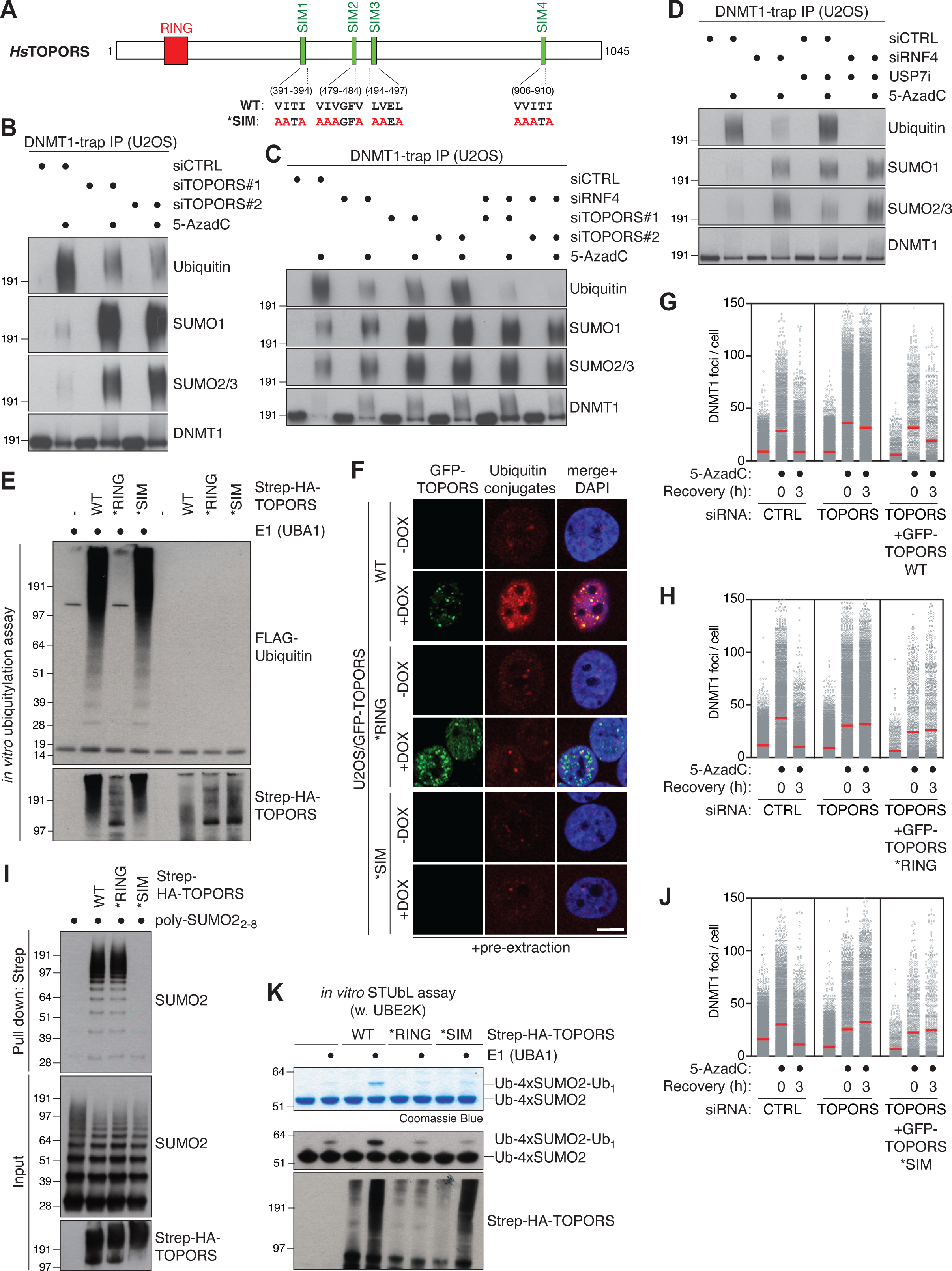
TOPORS functions as a SUMO-targeted ubiquitin ligase in DPC repair. **A.** Domain organization of human TOPORS, showing location of the RING domain and four predicted SIMs, as well as amino acid substitutions introduced to generate the TOPORS *SIM mutant. **B.** Immunoblot analysis of U2OS cells transfected with non-targeting control (CTRL) or TOPORS siRNAs for 72 h, treated or not with 5-AzadC for 30 min, collected and subjected to DNMT1 IP under denaturing conditions. **C.** As in (B), except that cells were treated with 5-AzadC 40 h after transfection with indicated siRNAs. **D.** As in (B), except that USP7i was added where indicated. **E.** Immunoblot analysis of *in vitro* ubiquitylation reactions containing recombinant Strep-HA-TOPORS proteins (**Figure S2C**) that were pre-incubated with Ub-VS and SUMO2-VS in reaction buffer for 10 min, supplemented with E1 (UBA1) and E2 (UBE2D1) enzymes, FLAG-ubiquitin and ATP and incubated at 30 °C for 10 min. **F.** Representative images of stable U2OS/GFP-TOPORS cell lines that were induced or not to express GFP-TOPORS proteins with Doxycycline (DOX), pre-extracted and fixed, and immunostained with antibody specific for ubiquitin conjugates (FK2). Scale bar, 10 μm. Representative images of non-pre-extracted U2OS/GFP-TOPORS cell lines in which the expression of the GFP-TOPORS *SIM mutant can be seen are shown in **Figure S2E**. **G.** U2OS cells were sequentially transfected with indicated siRNAs and expression plasmids (GFP-TOPORS WT and mCherry-H2B). Cells were then subjected to a single-round thymidine synchronization in early S phase and, upon release from the block, treated with 5-AzadC for 30 min, collected and pre-extracted at the indicated times. After DNMT1 immunostaining, transfected cells were gated based on mCherry positivity and DNMT1 foci formation in gated cells was analyzed by QIBC (red bars, mean; >1,030 cells analyzed per condition). Data are representative of 3 independent experiments. **H.** As in (G), except that cells were transfected with GFP-TOPORS *RING and mCherry-H2B plasmids (red bars, mean; >680 cells analyzed per condition). Data are representative of 3 independent experiments. **I.** Immunoblot analysis of recombinant Strep-HA-TOPORS proteins that were incubated with poly-SUMO2 chains and subjected to Strep-Tactin pulldown. **J.** As in (G), except that cells were transfected with GFP-TOPORS *SIM and mCherry-H2B plasmids (red bars, mean; >910 cells analyzed per condition). Data are representative of 3 independent experiments. **K.** Coomassie staining (top) and immunoblot analysis using SUMO2 and HA antibodies (bottom) of *in vitro* STUbL reactions containing recombinant Strep-HA-TOPORS proteins that were pre-incubated with Ub-VS and SUMO2-VS in reaction buffer for 10 min, supplemented with E1 (UBA1) and E2 (UBE2D1) enzymes, FLAG-ubiquitin, 4xSUMO2 STUbL substrate and ATP and incubated at 37 °C for 2 h.

This suggests that RNF4 and TOPORS both contribute to DNMT1 DPC ubiquitylation and together are responsible for the bulk of these modifications. Quantitative proteomic analysis further indicated an inverse impact of both TOPORS and RNF4 knockdown on 5-AzadC-induced DNMT1 ubiquitylation and SUMOylation levels (**Figure S2B; Table S2**), suggesting an uncoupling of these modifications. Similar to the effect of TOPORS knockdown, treatment with USP7i that rapidly depletes the endogenous TOPORS pool (**Figure 1H**) exacerbated the DNMT1 DPC ubiquitylation defect in cells lacking RNF4 even though USP7i exposure alone led to a moderate increase in DNMT1 ubiquitylation, possibly reflecting its previously described role in deubiquitylating SUMO and/or stabilizing DNMT1 itself ^4,30–32^ (**Figure 2D**). These observations suggested that TOPORS might act as an E3 ligase catalyzing ubiquitylation, but not SUMOylation, of DNMT1 DPCs in parallel with RNF4. In line with this, purified recombinant wild-type (WT) TOPORS was highly active as an E3 ubiquitin ligase *in vitro*, and this activity was fully abrogated by introduction of point mutations predicted to disrupt the E2-binding capacity of its RING domain (*RING) (**Figure 2E; Figure S2C**). By contrast, little SUMO E3 ligase activity of TOPORS was apparent *in vitro*, and no difference between TOPORS WT and *RING was observed (**Figure S2D**). In line with this, using cell lines conditionally expressing GFP-TOPORS proteins (**Figure S2E**), we noted that the detergent-resistant, chromatin-associated nuclear foci formed by GFP-TOPORS displayed strong enrichment of ubiquitin conjugates in a RING-dependent manner, whereas both GFP-TOPORS WT and *RING foci colocalized with, and showed a similar level of, SUMO2/3 (**Figure 2F; Figure S2F**). Ectopically expressed GFP-TOPORS WT but not *RING alleviated the DNMT1 DPC resolution defect resulting from depletion of endogenous TOPORS (**Figure 2G,H**), indicating that TOPORS E3 ligase activity via the RING domain is required for DNMT1 DPC resolution. In fact, overexpression of catalytically inactive TOPORS not only failed to rescue the DNMT1 DPC repair defect in cells lacking endogenous TOPORS but had a dominant-negative impact on DNMT1 DPC resolution in the presence of endogenous TOPORS (**Figure S2G**). Collectively, these data suggest that TOPORS acts as an E3 ubiquitin ligase for DNMT1 DPCs.

Given the similar behavior and requirements of RNF4 and TOPORS in SUMO-dependent DPC resolution, we considered the possibility that TOPORS might function as a STUbL. In support of this idea, TOPORS interacted strongly with poly-SUMO chains *in vitro* (**Figure 2I**), as also indicated by global proteomics-based profiling of SUMO-binding proteins in human cells ^33^. Inspection of the primary sequence of TOPORS revealed four prospective conserved SIMs in its middle and C-terminal portions (**Figure 2A**). Simultaneous functional disruption of these motifs by introduction of alanine substitutions in their hydrophobic core (*SIM; **Figure 2A**) had no impact on TOPORS E3 ubiquitin ligase activity *in vitro* (**Figure 2E**) but abrogated TOPORS interaction with SUMO2 chains and accumulation in detergent-resistant nuclear SUMO2/3-positive foci (**Figure 2F**; compare with **Figure S2F**; **Figure 2I**). Consistent with the presence of multiple SIMs, TOPORS displayed a strong preference for binding to long poly-SUMO2 chains and showed no detectable interaction with monomeric SUMO2 (**Figure 2I; Figure S2H**). Despite it maintained full E3 ubiquitin ligase activity *in vitro*, ectopic expression of the TOPORS *SIM mutant did not promote increased nuclear ubiquitylation unlike GFP-TOPORS WT (**Figure 2F**), suggesting that TOPORS might function as a STUbL. Moreover, ectopic TOPORS *SIM expression failed to rescue the DNMT1 DPC repair defect of cells depleted for endogenous TOPORS and did not interact with DNMT1 DPC sites, unlike WT TOPORS (**Figure 2G,J; Figure S2I**). To directly assay for TOPORS STUbL activity, further anticipated based on the combination of ubiquitin E3 ligase activity and SIMs characteristic of known STUbLs, we performed *in vitro* ubiquitylation assays with TOPORS proteins purified from HEK293T cells and linearly fused 4xSUMO2 proteins mimicking a poly-SUMO2 chain ^34^. Consistent with TOPORS acting as a *bona fide* STUbL, we found that it stimulated ubiquitylation of 4xSUMO2 substrates in a RING- and SIM-dependent manner (**Figure 2K; Figure S2J**). However, TOPORS was much less efficient than bacterially purified RNF4 in promoting 4xSUMO2 ubiquitylation in the presence of the generic E2 ubiquitin-conjugating enzyme UBE2D1 that supports the ubiquitin ligase activity of many E3s *in vitro* (**Figure S2J**), raising the possibility that TOPORS may be more active as a STUbL in the presence of other, specific E2s. Because the E2 enzyme UBE2K was among the hits in our genetic screen for factors required for DNMT1 DPC repair (**Figure 1A,B; Table S1**), we considered the possibility that it might function with TOPORS. Indeed, TOPORS displayed markedly higher STUbL activity with UBE2K than with UBE2D1, comparable to RNF4 whose 4xSUMO2-directed E3 ligase activity was supported to a similar extent by UBE2D1 and UBE2K (**Figure S2J**). However, TOPORS and RNF4 only stimulated UBE2K-dependent ubiquitylation of a ubiquitin-fused 4xSUMO2 protein (Ub-4xSUMO2) but not 4xSUMO2 *per se* (**Figure 2K; Figure S2J**), in agreement with the known role of UBE2K as an E2 that elongates ubiquitin modifications by catalyzing their K48 linkage-specific ubiquitylation ^35–37^. The preference of TOPORS for UBE2K suggests that it could function as a STUbL that is important for extending pre-existing ubiquitin modifications on SUMOylated substrates, e.g. those generated by RNF4, into K48-linked chains. In line with this and our screen data, UBE2K knockdown delayed the resolution of DNMT1 DPCs (**Figure S2K,L**). We conclude from these findings that human TOPORS is a novel STUbL that promotes SUMO-dependent DPC repair by means of this activity in conjunction with RNF4.

### TOPORS and RNF4 cooperatively drive multiple STUbL-driven responses

The concerted action of RNF4- and TOPORS-dependent STUbL activities in promoting SUMO-dependent DPC resolution raised the question of whether this partnership extends to other cellular STUbL-driven processes. We therefore used an established *PML* knockout (KO) cell line stably reconstituted with YFP-tagged PML ^38^ to assess whether TOPORS is involved in arsenic-induced PML degradation, which like DNA replication-independent DPC resolution is mediated by SUMOylation and RNF4 STUbL activity ^8,9^. Interestingly, paralleling the impact of RNF4 depletion (**Figure S3A**), knockdown of TOPORS diminished arsenic-induced PML ubiquitylation whereas SUMO-modified forms of PML accumulated (**Figure 3A**). This effect appeared particularly prominent for SUMO1, a trend that was also seen for DNMT1 DPCs (**Figure 3A**; **Figure 2B; Figure S2A**). Consistently, TOPORS knockdown strongly impaired PML body degradation upon arsenic treatment, to a greater extent than the defect seen in RNF4-depleted cells (**Figure 3B; Figure S3B**). Inhibition of USP7 activity likewise impaired the timely clearance of PML bodies after arsenic treatment (**Figure 3C**). While individual depletion of TOPORS or RNF4 for a shorter (40-h) period only caused a mild reduction in arsenic-induced PML ubiquitylation, their combined knockdown largely abrogated this modification (**Figure 3D**). This suggests that RNF4 and TOPORS cooperatively promote PML ubiquitylation and subsequent degradation upon arsenic treatment, mirroring their joint involvement in DPC resolution. Further linking TOPORS to PML body degradation, proteomic analysis of YFP-PML pulldowns showed increased association of endogenous TOPORS with PML upon arsenic treatment (**Figure 3E; Figure S3C; Table S3**) ^38^. Moreover, ectopic expression of E3 ligase-deficient TOPORS, but not the WT protein, significantly stabilized PML bodies (**Figure S3D**), indicating that TOPORS E3 ligase activity is important for turning over these structures. In addition to DPC and PML degradation, RNF4 and SUMOylation have been implicated in the proteasome-dependent clearance of defective ribosomal products (DRiPs) arising from treatment with the translational inhibitor O-Propargyl-puromycin (OP-puro) ^39–41^. We observed that both RNF4 knockdown and inhibitors of SUMOylation, ubiquitylation and the proteasome delayed the resolution of OP-puro-labeled DRiPs (**Figure 3F; Figure S3E**), supporting an important role of STUbL activity in this process. Depletion of TOPORS by independent siRNAs likewise delayed the clearance of DRiPs (**Figure 3F**), suggesting that TOPORS and RNF4 jointly promote multiple STUbL-driven pathways. In line with this notion, we noted that co-depletion of RNF4 and TOPORS, but not individual knockdown of either ligase, led to a striking accumulation of very high molecular weight SUMOylated proteins in total cell extracts manifesting as slow-migrating ‘well products’ in SDS-PAGE (**Figure 3G**), suggesting a severe defect in processing SUMO-modified proteins when both RNF4 and TOPORS are absent in otherwise unstressed cells. As seen for other phenotypes, USP7i treatment recapitulated the impact of TOPORS depletion in causing accumulation of hyper-SUMOylated proteins in cells lacking RNF4 but not in control cells (**Figure 3H**). Together, these findings suggest that the coupling of TOPORS and RNF4 E3 ubiquitin ligase activities is a general principle in STUbL-driven cellular pathways, exemplified by their joint action in SUMO-dependent processing of DPCs, PML bodies and DRiPs.

**Figure 3.**
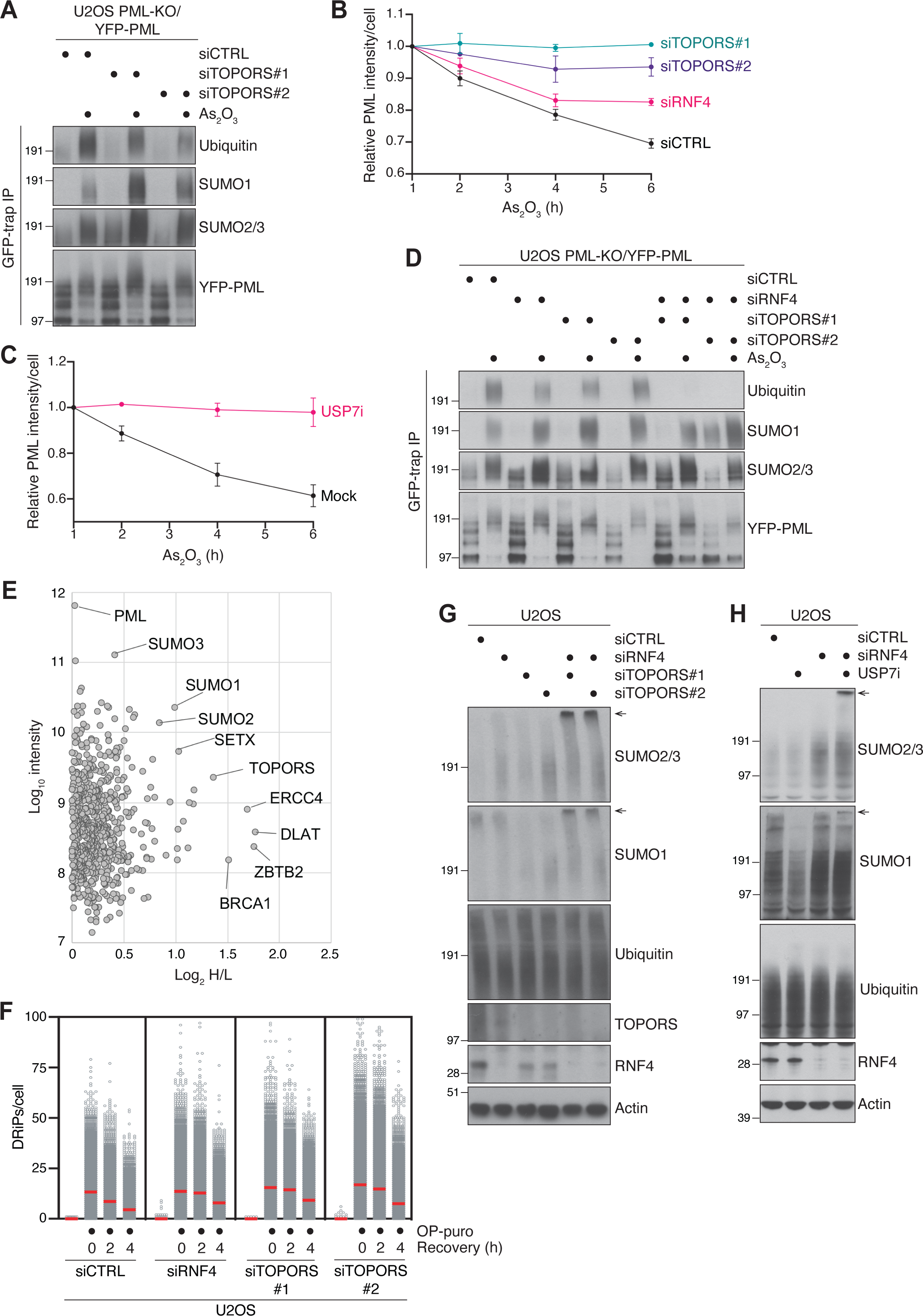
TOPORS and RNF4 cooperatively drive multiple STUbL-driven responses. **A.** U2OS PML-KO cells stably reconstituted with YFP-tagged PML-V (U2OS PML-KO/YFP-PML) were transfected with indicated siRNAs for 72 h, exposed to arsenic (As_2_O_3_) and collected 1 h later. Cells were subjected to GFP IP under denaturing conditions followed by immunoblotting with indicated antibodies. **B.** U2OS PML-KO/YFP-PML cells transfected with indicated siRNAs were treated with arsenic and fixed at indicated time points. YFP-PML intensity was analyzed by quantitative image-based cytometry (QIBC) (mean±SEM; *n*=3 independent experiments). **C.** U2OS PML-KO/YFP-PML cells were exposed to arsenic in the presence or absence of USP7i, fixed at the indicated times and analyzed as in (B) (mean±SEM; *n*=3 independent experiments). **D.** As in (A), except that cells were treated with arsenic 40 h after transfection with indicated siRNAs. **E.** U2OS PML-KO/YFP-PML cells were grown in SILAC medium and exposed to 1 µM arsenic for 2 h (Heavy condition) or left untreated (Light condition). PML bodies were enriched by affinity purification using GFP nanobody crosslinked to magnetic beads, and associated proteins identified and quantified by mass spectrometry. Log2 of protein abundance ratios are displayed on the x-axis with log10 protein intensity values on the y-axis. Selected outliers with increased abundance and PML itself are indicated. **F.** U2OS cells transfected with indicated siRNAs were treated with MG132 and O-Propargyl-puromycin (OP-puro) for 4 h, washed and released into fresh media, and collected at the indicated times. OP-puro was click-conjugated to fluorescent azide and analyzed by QIBC (red bars, mean; >7,700 cells analyzed per condition). Data are representative of 3 independent experiments. **G.** Immunoblot analysis of U2OS cells transfected with indicated siRNAs for 40 h. Slow-migrating, hyper-SUMOylated proteins (‘well products’) are indicated by arrows. **H.** Immunoblot analysis of U2OS cells transfected with indicated siRNAs for 48 h and grown in the absence or presence of USP7i for an additional 12 h. Slow-migrating, hyper-SUMOylated proteins (‘well products’) are indicated by arrows.

### TOPORS and RNF4 generate complex ubiquitin topologies on STUbL substrates to promote p97 recruitment

We next asked why both RNF4 and TOPORS are necessary for efficient STUbL substrate turnover via seemingly similar mechanisms. Recent work, which we independently verified (**Figure S4A,B**), demonstrated that p97 activity is essential for SUMO-dependent degradation of both DNMT1 DPCs and PML bodies ^16,38^. Strikingly, indicative of a role for TOPORS STUbL activity in promoting p97 recruitment to SUMOylated proteins, we noted that the nuclear foci formed by stably expressed GFP-TOPORS not only accumulated high levels of ubiquitin conjugates but also displayed strong positivity for p97, in a manner that was fully dependent on TOPORS E3 ligase activity and SUMO binding (**Figure 2F**; **Figure 4A**). Consistently, knockdown of TOPORS markedly reduced p97 recruitment to DNMT1 DPC sites, to an extent that was similar to the impact of RNF4 depletion (**Figure 4B**). Moreover, unlike WT TOPORS, cells expressing the TOPORS *RING or *SIM mutants failed to support p97 recruitment to DNMT1 DPC sites (**Figure S4C**). Depletion of TOPORS or RNF4 likewise diminished the occupancy of p97 in PML bodies upon arsenic treatment and p97 association with DRiPs (**Figure 4C,D**). Furthermore, we found that inhibition of p97 activity phenocopied the accumulation of hyper-SUMOylated proteins seen upon combined RNF4 and TOPORS loss (**Figure S4D**), suggesting that these STUbLs together ensure efficient processing of SUMOylated proteins by promoting p97 recruitment. However, the role of RNF4 and TOPORS in promoting p97-dependent processing of ubiquitylated substrates may be specific for STUbL targets, as neither RNF4 nor TOPORS depletion affected the levels of a ubiquitylated model substrate (Ub(G76V)-GFP) whose proteasomal turnover requires p97 activity ^42^, but not SUMOylation (**Figure S4E**). These findings suggested that both TOPORS and RNF4 make important contributions to the generation of ubiquitin signals underlying efficient p97 recruitment to STUbL substrates. To explore this possibility directly, we quantified the abundance of individual ubiquitin linkages in our proteomic analysis of DNMT1 DPCs isolated under stringent denaturing conditions (**Figure S2A**). This showed that DNMT1 DPCs are mainly modified by K48-linked ubiquitin chains (**Figure S4F; Table S2**), a major signal for p97- and proteasome-dependent turnover ^43^, and that both TOPORS and RNF4 depletion leads to a sizeable decrease in these modifications (**Figure 4E; Table S2**), in accordance with our findings above. By contrast, RNF4 but not TOPORS was required for the comparatively lower level of K63- and K11-linked ubiquitylation of DNMT1 that was also induced by 5-AzadC treatment (**Figure 4E; Figure S4F; Table S2**). Similarly, knockdown of RNF4, but not TOPORS, reduced the extent of arsenic-induced K63-linked ubiquitin chain formation on PML (**Figure S4G**). Interestingly, probing total ubiquitylated cellular proteins isolated by the Multi-Dsk affinity reagent ^44^ under denaturing conditions to prevent co-purification of associated factors revealed that loss of TOPORS, but not RNF4, led to a drastic decline in proteins co-modified by ubiquitin and SUMO1, whereas levels of ubiquitylated proteins modified by SUMO2/3 were moderately increased (**Figure 4F**). This suggests that TOPORS may provide a principal cellular source of STUbL activity towards SUMO1-modified proteins, in line with the more pronounced impact of TOPORS loss on DNMT1 DPC and PML modification by SUMO1 than SUMO2/3 (**Figure 2B**; **Figure 3A; Figure S2A**). Together, these data suggest that TOPORS and RNF4 have both overlapping and complementary roles in ubiquitylating STUbL substrates, with both E3s contributing to their K48-linked ubiquitylation while TOPORS is more important for ubiquitylation of SUMO1-modified proteins and RNF4 is the primary driver of non-K48-linked ubiquitylation of SUMO targets. The combined action of TOPORS and RNF4 STUbL activities may thus enable the generation of complex ubiquitin topologies on both SUMO1- and SUMO2/3-modified proteins to provide optimal recruitment signals for p97-cofactor complexes, facilitating proteolytic processing of unfolded STUbL substrates by the proteasome.

**Figure 4.**
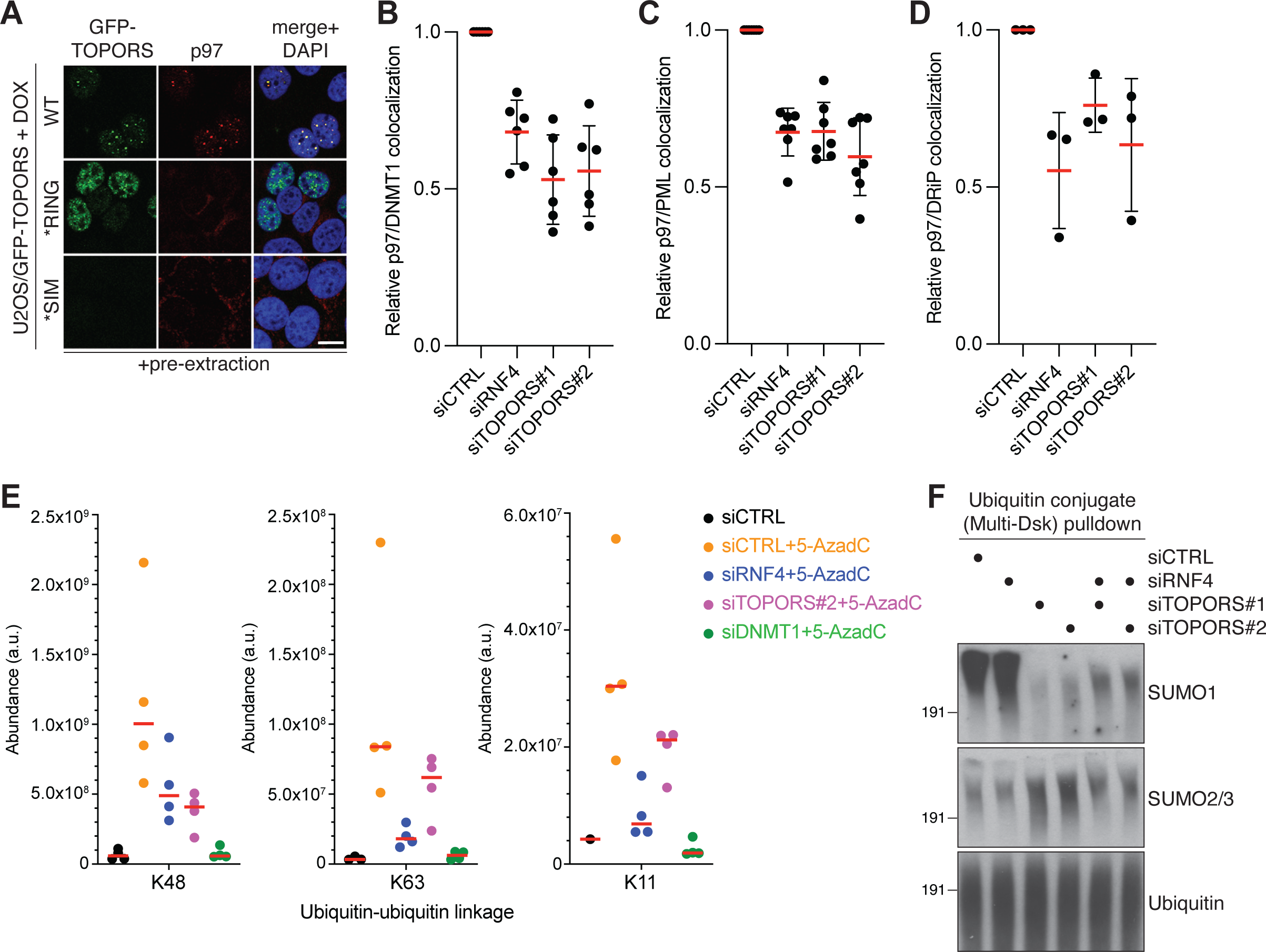
TOPORS and RNF4 generate complex ubiquitin topologies on STUbL substrates to promote p97 recruitment. **A.** Representative images of DOX-treated U2OS/GFP-TOPORS cell lines immunostained with p97 antibody after pre-extraction and fixation. Scale bar, 10 μm. **B.** U2OS cells transfected with indicated siRNAs were released from single thymidine synchronization in early S phase and treated with 5-AzadC for 30 min. Cells were collected, pre-extracted and immunostained with p97 and DNMT1 antibodies. Intensity of p97 and DNMT1 in DNMT1 DPC foci was analyzed by QIBC (mean±SEM; *n*=6 independent experiments). **C.** U2OS PML-KO/YFP-PML cells transfected with indicated siRNAs for 72 h and treated with arsenic for 1 h were collected, pre-extracted and immunostained with p97 antibody. Intensity of p97 and YFP in YFP-PML bodies was analyzed by QIBC (mean±SEM; *n*=7 independent experiments). **D.** U2OS cells transfected with indicated siRNAs were treated with MG132 and O-Propargyl-puromycin (OP-puro) for 4 h. Cells were collected, pre-extracted and immunostained with p97 antibody. OP-puro was click-conjugated to fluorescent azide, and p97 and OP-puro intensity in defective ribosomal products (DRiPs) was analyzed by QIBC (mean±SEM; *n*=3 independent experiments). **E.** Mass spectrometry (MS)-based analysis of ubiquitin-ubiquitin linkages on DNMT1. U2OS cells transfected with indicated siRNAs were subjected to DNMT1 IP under denaturing conditions to isolate DNMT1 but not associated proteins. Samples were digested with trypsin, after which peptides were purified and MS was used to identify di-glycine remnants (*n*=4 independent experiments). **F.** Immunoblot analysis of U2OS cells transfected with indicated siRNAs for 40 h and subjected to Multi-Dsk pulldown under denaturing conditions to isolate total cellular ubiquitin conjugates but not associated, non-covalently bound proteins. Input blots are shown in Figure 3G.

### Combined loss of TOPORS and RNF4 is synthetic lethal

To further interrogate the STUbL functions of RNF4 and TOPORS and their interrelationship, we screened for genetic interactions affecting cell fitness using HAP1 cell lines lacking TOPORS or RNF4 function (**Figure S1C; Figure S5A,B**). Due to concerns about possible long-term growth defects associated with full KO of RNF4 but not TOPORS that were apparent from genetic screens in unchallenged HAP1 WT cells (**Figure 5A**) ^45^, we endogenously tagged RNF4 with a C-terminal degron (FKBP12(F36V)-HA; dTAG-HA) to enable its efficient conditional depletion upon addition of the dTAG-13 degrader (**Figure S5B**). The functionality of TOPORS KO or dTAG-13-dependent RNF4 depletion in these cell lines was validated by their defective ability to degrade DNMT1 in S phase upon 5-AzadC treatment, contrary to WT cells (**Figure 1C; Figure S5C**). Intriguingly, fitness-based gene-trap mutagenesis screens in TOPORS-deficient cells identified *RNF4* as a major synthetic lethal hit (**Figure 5A-D; Table S4**). Correspondingly, disruption of *TOPORS* gave rise to a strong synthetic lethality effect in RNF4-deficient cells (**Figure 5A-D; Table S4**). Underscoring the critical role of USP7 activity in sustaining TOPORS expression, *USP7* was also a strong synthetic lethality hit in cells lacking RNF4, but not in TOPORS-KO cells (**Figure 5A-D**). We validated a profound loss of proliferative potential resulting from co-depletion of TOPORS and RNF4 in U2OS cells (**Figure 5E; Figure S5D**). We also confirmed that low dose treatment with USP7i strongly suppressed the proliferation of RNF4-deficient cells but had little impact in parental and TOPORS-depleted cells (**Figure S5E-G**). Co-depletion of TOPORS and RNF4, but not individual knockdown of either ligase, led to a prominent increase in levels of PARP1 cleavage, a marker of apoptosis (**Figure 5F**), suggesting that the joint loss of TOPORS and RNF4 is incompatible with cell survival. USP7i addition to RNF4-depleted but not control cells produced a similar effect (**Figure S5H**). These findings demonstrate that the combined actions of TOPORS and RNF4 are essential for cell viability and proliferation, corroborating the critical importance of concerted TOPORS and RNF4 STUbL activities in promoting cellular stress responses and the turnover of SUMOylated proteins.

**Figure 5.**
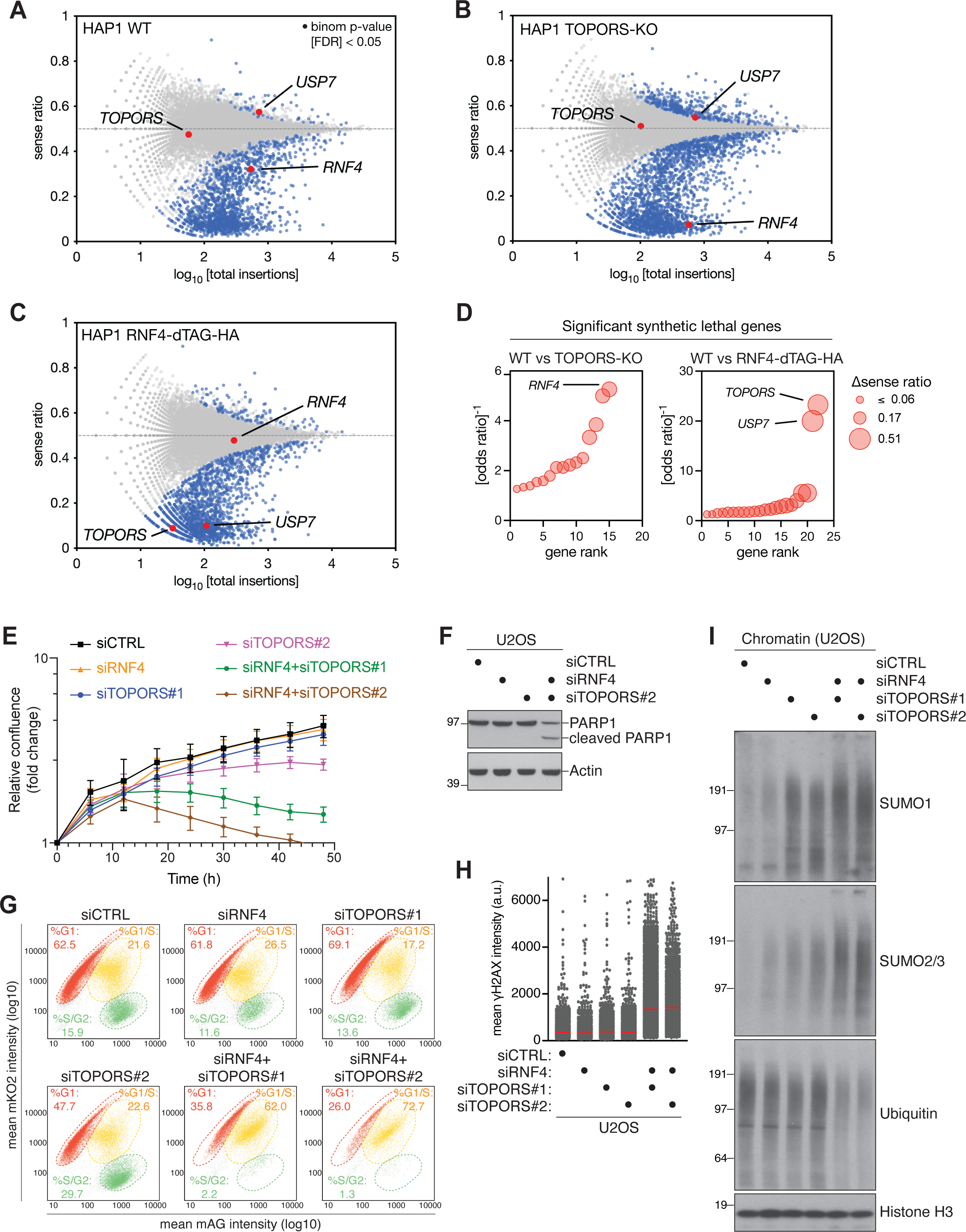
Combined loss of TOPORS and RNF4 is synthetic lethal. **A.** Haploid genetic fitness screen in HAP1 WT cells, presented as a fishtail plot in which genes are plotted according to ratio of sense/anti-sense orientation of gene-trap insertions (y-axis) and the total number of insertions in a particular gene (x-axis) (*n*=4 biological replicates (independent clones)). Blue dots represent genes with significant sense bias (binom p-value<0.05 FDR corrected, across all replicates). A single representative replicate is shown. **B.** As in (A), but for HAP1 TOPORS-KO cells (*n*=2 biological replicates (independent clones)). **C.** As in (A), but for HAP1 RNF4-dTAG-HA cells treated with 0.25 µM dTAG13 for 10 days (*n*=2 biological replicates (independent clones)). **D.** Gene-rank plot for significant synthetic lethal genes for HAP1 TOPORS-KO (left) and RNF4-dTAG-HA (right) compared to HAP1 WT (Fisher’s exact test; p<0.05; odds ratio<0.8). The size of the dot represents the difference (Δ) in sense ratio bias. **E.** Normalized logarithmic proliferation quantification for U2OS cells transfected with indicated siRNAs for the indicated times, as determined by Incucyte image-based confluence analysis (mean±SEM; *n*=3 independent experiments). Imaging started at 24 h after siRNA transfection. **F.** Immunoblot analysis of U2OS cells transfected with indicated siRNAs for 48 h. **G.** U2OS Fucci cells were transfected with indicated siRNAs for 36 h. To determine cell cycle position, CDT1 (mKO2-hCdt1+) and Geminin (mAG-hGem+) intensities were analyzed by QIBC. Quantification of cells in different cell cycle phases was done using the indicated gates. Data are representative of two independent experiments. **H.** U2OS cells were transfected with indicated siRNAs, treated with thymidine for 18 h and released into S phase for 6 h. Cells were immunostained with ψH2AX antibody and analyzed by QIBC (red bars, mean; >3,900 cells analyzed per condition). Data are representative of two independent experiments. **I.** Immunoblot analysis of chromatin-enriched fractions of U2OS cells transfected with indicated siRNAs for 40 h.

To understand how the simultaneous loss of TOPORS and RNF4 undermines cell proliferation, we performed live-cell imaging experiments using a Fucci reporter cell line, in which individual cell cycle phases and cell cycle progression can be visualized and tracked in living cells via fluorescent reporters ^46^. We observed extensive cell death upon co-depletion of TOPORS and RNF4 as expected, and this predominantly occurred around early S phase (**Figure S6A**). In line with this, quantification of Fucci cells in different cell cycle phases revealed a strong accumulation of cells at the G1/S transition and a concomitant loss of S/G2 phase cells when both TOPORS and RNF4, but not either ligase alone, were depleted (**Figure 5G**). Again, this effect was phenocopied by USP7i treatment of RNF4-depleted cells (**Figure S6B**). These observations suggested that combined loss of RNF4 and TOPORS gives rise to a severe DNA replication defect. To test this prediction, we performed EdU labeling experiments with cells released from G1/S transition arrest by a double thymidine block. Individual depletion of TOPORS or RNF4 had little impact on EdU incorporation efficiency (**Figure S6C**). By contrast, cells lacking both TOPORS and RNF4 displayed markedly reduced DNA synthesis rates in early S phase and failed to sustain replication further into S phase (**Figure S6C**). RNF4-depleted cells exposed to low dose USP7i treatment likewise exhibited diminished EdU incorporation rates (**Figure S6D**). The DNA replication defect arising from combined TOPORS and RNF4 knockdown coincided with accumulation of DNA damage as evidenced by an increased level of γH2AX formation in S phase cells (**Figure 5H**), suggesting that simultaneous loss of TOPORS and RNF4 leads to extensive replication fork stalling and/or collapse. We surmised that this DNA replication defect in cells co-depleted for RNF4 and TOPORS might be caused by obstacles to replisome progression in the form of SUMOylated proteins that fail to get extracted from the chromatin template, in keeping with the known role of STUbL activity in this process ^3^. Indeed, combined knockdown of TOPORS and RNF4 led to a strong accumulation of SUMOylated proteins on chromatin and a concomitant decrease in ubiquitylation, with TOPORS depletion alone being sufficient to trigger a substantial increase in SUMO1-modified chromatin-bound proteins (**Figure 5I**). Collectively, these data suggest that the combined STUbL activities of TOPORS and RNF4 are not only instrumental for SUMO-dependent stress responses but also exert an essential role during a normal cell cycle by ubiquitylating and turning over SUMOylated proteins from chromatin to allow unhindered advancement of the replication machinery.

## Discussion

By catalyzing selective ubiquitylation of SUMOylated proteins, STUbLs enable direct crosstalk between ubiquitin- and SUMO-mediated signaling in cells and act as key effectors in a range of stress responses, whose pharmacological modulation has shown strong potential in cancer treatment. A detailed understanding of the mechanisms and functions of STUbLs in cell biology is therefore of considerable importance. In this study, we discovered that the human TOPORS protein is a novel STUbL by virtue of its RING domain and a poly-SUMO-binding SIM cluster, a configuration analogous to that of other known STUbLs such as RNF4 and RNF111. The precise enzymatic function of TOPORS has been uncertain, as previous studies provided evidence for TOPORS functioning as an E3 ligase for both ubiquitin and SUMO ^25–29^. While we do not exclude that TOPORS may serve as a SUMO E3 ligase in some contexts, our data from *in vitro* and cell-based experiments show that TOPORS is by far more active as an E3 ubiquitin ligase by means of the RING domain, and they collectively favor the notion that the ubiquitin ligase function of TOPORS provides the critical activity for its role in STUbL-dependent processes.

Our findings show that TOPORS makes key contributions to established RNF4-driven pathways, including SUMO-dependent DPC resolution and PML body degradation. Rather than simply serving as a backup to RNF4, our data suggest that TOPORS and RNF4 are both required for the efficiency of STUbL reactions. In fact, in our experiments TOPORS depletion consistently gave rise to stronger DPC repair and PML body processing defects than RNF4 knockdown. Importantly, the dedicated partnership between TOPORS and RNF4 not only underpins the efficacy of SUMO-driven stress responses but is also essential for cell proliferation and viability under basal, unstressed conditions. Accordingly, in the absence of both TOPORS and RNF4, but not either STUbL alone, cells accrue high levels of hyper-SUMOylated proteins, presumably due to a failure to process these species via the p97-proteasome pathway resulting from their lack of ubiquitylation. This is accompanied by a strong DNA replication defect that may be a direct consequence of the inability to properly displace SUMOylated proteins from the chromatin template, obstructing the progression of the replication machinery. Indeed, we found that combined loss of TOPORS and RNF4 disrupts the normal balance of SUMO and ubiquitin modifications on chromatin, which has been proposed to be critical for maintaining the activity of replication forks ^47^. Given that STUbL activity is important for removing both DPCs and non-covalently bound proteins from DNA ^3,14,15^, we consider it likely that SUMOylated forms of both classes of proteins accumulate on chromatin and contribute to impeding replication when both TOPORS and RNF4 are absent. Our findings thus reveal an unexpected mechanistic complexity of major STUbL-driven processes, suggesting that the concerted action of TOPORS and RNF4 is a general modus operandi of these pathways with an indispensable role in cell proliferation.

A key question emanating from this discovery is why both TOPORS and RNF4 are needed for the productive execution of STUbL-mediated responses. Together with our other findings, the notion that combined RNF4 and TOPORS loss is lethal while their individual depletion is not suggests that these STUbLs have both overlapping and non-redundant roles in ubiquitylating SUMOylated proteins. According to this scenario, STUbL-driven pathways become inefficient but retain basic functionality in the absence of either ligase, whereas loss of both TOPORS and RNF4 leads to a profound defect in the ubiquitylation and proteolytic processing of SUMOylated proteins. This is in good agreement with our finding that both TOPORS and RNF4 make sizeable contributions to the modification of SUMOylated DNMT1 DPCs by K48-linked ubiquitin chains. At the same time, RNF4 appears to be the main effector of non-K48-linked ubiquitylation of STUbL substrates, consistent with its ability to catalyze K63-linked ubiquitin chain formation ^48^, whereas our data suggest that TOPORS is particularly important for the ubiquitylation of SUMO1-modified proteins. Such a partial division of labor may allow for the efficient generation of a complex ubiquitin landscape on SUMO substrates to optimally support p97 recruitment, which we found requires both TOPORS and RNF4 function. K48-linked ubiquitin chains provide a primary recruitment signal for p97-cofactor complexes ^43^, in line with our finding that K48-linkages account for the majority of DNMT1 DPC-associated ubiquitin conjugates. The observation that TOPORS has little activity in the presence of the generic E2 enzyme UBE2D1 but is considerably more active *in vitro* with UBE2K, an E2 ubiquitin-conjugating enzyme specialized in extending pre-existing ubiquitin modifications into K48-linked chains, is well aligned with our proteomic data suggesting that TOPORS is mainly, if not exclusively, important for K48-linked ubiquitylation of DNMT1 DPCs. Intriguingly, recent work ^37,49^ showed that UBE2K is particularly efficient at promoting K48-linked ubiquitylation of K63-linked ubiquitin chains, leading to the generation of K48-K63 branched ubiquitin signals. Moreover, a very recent study suggested that K48-K63 branched ubiquitin chains may be a particularly strong recognition signal for p97 ^50^. These insights suggest a model for the requirement of concerted TOPORS and RNF4 STUbL activities wherein TOPORS-UBE2K promotes K48-linked ubiquitylation of RNF4-generated K63 linkages on STUbL substrates to facilitate their efficient p97- and proteasome-dependent turnover. Consistently, we found that UBE2K is rate-limiting for timely DNMT1 DPC resolution.

In addition to making differential contributions to the ubiquitin landscape formed on SUMOylated target proteins, it is possible that TOPORS and RNF4 differ with respect to the poly-SUMO chain topologies they preferentially recognize, in keeping with the different spacing of their multiple SIMs, as is also the case for RNF4 and RNF111 ^5^. In this regard, it is noteworthy that TOPORS appears to be particularly important for ubiquitylation of SUMO1-modified proteins in cells. It has been shown that poly-SUMO2/3 chain binding by the closely spaced tandem SIMs in RNF4 converts inactive monomeric RNF4 into an active dimer ^6^, and it is conceivable that such RNF4 activation could be difficult to achieve by multiple mono-SUMO1 modifications on target proteins, unless they are in close proximity. With the more scattered distribution of its SIMs, TOPORS might be better configured to recognize and ubiquitylate proteins modified by multi-mono-SUMO1 modifications. Indeed, we found that TOPORS knockdown led to an accumulation of SUMO1 modifications on DNMT1 DPCs, PML and chromatin-associated proteins, while the corresponding effects for SUMO2/3 were less pronounced. Interestingly, we showed in recent work that a patient-associated PML L217F mutant that is refractory to arsenic-induced degradation exhibited normal levels of SUMO2/3 modification and RNF4 recruitment but markedly reduced SUMO1 conjugation^51^. It is tempting to speculate that this PML mutant fails to undergo degradation upon arsenic treatment due to impaired ubiquitylation by TOPORS. Addressing the precise relationships between individual STUbLs and different SUMO chain types will be important but is challenged by the limited current insights into these polymers, their low abundance in cells and a shortage of suitable tools and methods for analyzing specific SUMO chain architectures ^1^. Finally, it is possible that the coordinated activities of TOPORS and RNF4 together enable the generation of ubiquitin chains on SUMO-modified proteins that are sufficiently long to facilitate p97 recruitment. Recent work revealed that the budding yeast p97 orthologue Cdc48 in complex with its core cofactor Ufd1-Npl4 requires a minimum chain length of around five ubiquitins to efficiently process ubiquitylated substrates ^52–54^, and the joint action of TOPORS and RNF4 E3 ligase activities on STUbL substrates may be important for surpassing this threshold. Irrespective of the precise mechanisms, it seems likely that the concerted actions of TOPORS and RNF4 enables flexibility in STUbL-driven processes targeting a multitude of cellular substrates, and that the relative importance of TOPORS and RNF4 STUbL activities may differ depending on the precise SUMOylated target and context.

Along with TOPORS, we found that USP7 deubiquitinase activity is crucial for sustaining the integrity of STUbL-dependent processes in conjunction with RNF4. Our collective evidence strongly suggests that this is exerted via a critical role of USP7 in underpinning TOPORS stability by antagonizing its auto-ubiquitylation. Recent system-wide proteomic studies indicated a highly selective loss of TOPORS expression upon brief treatment with USP7 inhibitors ^22,23^, and we verified that USP7i leads to a precipitous decline in TOPORS expression manifesting fully already within 10 min of treatment. Consistently, USP7i addition to RNF4-depleted cells recapitulated major phenotypes resulting from simultaneous loss of TOPORS and RNF4 but did not produce a similar effect when administered to cells lacking TOPORS. Although USP7 has been reported to function as a SUMO-targeted ubiquitin protease that antagonizes SUMO-dependent protein ubiquitylation and degradation ^4^, our data argue against such an activity impacting STUbL substrates such as DNMT1 DPCs directly as being a main determinant for the role of USP7 in promoting RNF4-mediated responses. In this regard, preventing USP7-mediated deubiquitylation of SUMOylated proteins to enhance their degradation would be expected to at least partially reverse, rather than exacerbate, the STUbL pathway defects caused by RNF4 depletion, which is not what we observed. Instead, the notion that TOPORS depletion and USP7 inhibition have virtually identical impacts on RNF4-deficient cells in multiple assays strongly suggests that at least in the context of STUbL-driven processes the key function of USP7 is to protect TOPORS from degradation.

In summary, our findings uncover TOPORS as a novel STUbL in human cells and establish that direct SUMO-ubiquitin crosstalk mediated by the concerted activities of TOPORS and RNF4 is essential for cell viability and proliferation. These improved insights into the mechanistic basis of major SUMO- and STUbL-dependent processes could have significant bearings on current, promising treatment strategies targeting the SUMO system in cancers and offer potential new opportunities for therapeutic intervention. Pharmacological modulation of STUbL-dependent processes with a key involvement of TOPORS including arsenic-induced PML body degradation and resolution of 5-AzadC-induced DPCs has documented clinical benefits ^19,20^. Moreover, many cancers are addicted to a hyper-active SUMO system as a means to cope with high levels of stress, and the specific SUMO E1 inhibitor TAK-981 has emerged as a promising anticancer agent, synergizing with immune checkpoint inhibitors in mice ^55^. Targeting the critical role of TOPORS in STUbL-driven cellular responses might provide a more specific and thus less toxic alternative to full inhibition of SUMOylation in some clinical contexts. In this regard, harnessing the notion that USP7 inhibitors, several of which display efficacy for cancer therapy in pre-clinical studies ^56^, potently deplete TOPORS and recapitulate the consequences of TOPORS loss may be particularly useful. Further studies of the mechanisms of TOPORS STUbL function and how this may be exploited therapeutically are therefore well warranted.

## Methods

### Cell culture

Human HeLa, U2OS, HAP1 and HEK293-6E cells were obtained from ATCC. HeLa, U2OS and HEK293-6E cells were propagated in Dulbecco’s modified Eagle’s medium (DMEM) supplemented with 10% (v/v) fetal bovine serum and 1% penicillin-streptomycin. Cells were cultured in humidified incubators at 37 °C with 5% CO_2_ and were regularly tested negative for mycoplasma infection. The cell lines were not authenticated. HAP1 cells were cultured in IMDM-medium (Gibco) supplemented with 10% heat-inactivated fetal calf serum (FCS; ThermoFisher Scientific) and penicillin-streptomycin-glutamine solution (Gibco). HeLa cells stably expressing GFP-DNMT1 (HeLa/GFP-DNMT1) were described previously ^11^. To generate inducible cell lines expressing WT and mutant GFP-TOPORS proteins, pcDNA5/FRT/TO/GFP/TOPORS plasmids were co-transfected with pOG44 into U2OS Flp-In T-Rex cells (ThermoFisher Scientific) using Lipofectamine3000 (Invitrogen). Clones were selected in medium containing Hygromycin B (ThermoFisher) and blasticidin (InvivoGen) and verified by microscopy.

Plasmid DNA transfections were performed using FuGENE 6 (Promega), Lipofectamine 2000 (Invitrogen) or Lipofectamine 3000 (Invitrogen), according to the manufacturers’ protocols. Cell cycle synchronizations were performed as described ^57^. Briefly, HeLa, U2OS or HAP1 cells were synchronized by single treatment with thymidine for 18 h. Unless otherwise indicated, the following doses of chemicals and genotoxic agents were used: 5-Aza-2’-deoxycytidine (5-AzadC; 10 μM, Sigma-Aldrich), 5-Ethynyl-2’-deoxyuridine (EdU; 20 µM, Sigma-Aldrich), arsenic trioxide (Arsenic; 1 µM, Sigma-Aldrich), formaldehyde (500 µM, Thermo Scientific Pierce), FT-671 (USP7i; 2 µM, MedChemExpress), MG132 (20 μM, Sigma-Aldrich), ML-792 (SUMOi; 2 μM, MedKoo Biosciences), MLN-7243 (Ub-E1i; 5 μM, Active Biochem), NMS-873 (p97i; 5 μM, Sigma-Aldrich), O-propargyl-puromycin (OP-puro; 25 µM, Invitrogen) and thymidine (2 mM, Sigma-Aldrich).

### Plasmids

Full length cDNAs encoding human TOPORS WT (codon optimized, Invitrogen GeneArt), *RING (I105A, L140A, K142A; codon optimized), and TOPORS *SIM (V391A, I392A, I394A, V479A, I480A, V481A, V484A, L494A, V495A, L497A, V906A, V907A, I908A, I910A; codon optimized) were synthesized by ThermoFisher Scientific, PCR-amplified and cloned into pcDNA5/FRT/TO/GFP via KpnI and NotI (for generation of stable cell lines) or pcDNA4/TO/Strep-HA via EcoRV and NotI (for protein expression).

### siRNAs

siRNA transfections were performed using Lipofectamine RNAiMAX (Invitrogen) according to the manufacturer’s instructions. All siRNAs were used at a final concentration of 20 nM unless otherwise indicated. The following siRNA oligonucleotides, whose knockdown efficiencies were validated where indicated, were used: non-targeting control (CTRL): 5’-GGGAUACCUAGACGUUCUA-3’; RNF4: 5’-GAAUGGACGUCUCAUCGUU-3’; TOPORS#1: 5’-GUCCUAAGGCCUUCGUAUAAU-3’; TOPORS#2: 5’-CCCUGCUCCUUCAUACGAA-3’; UBE2K#1: 5’-GAAUCAAGCGGGAGUUCAA-3’; UBE2K#2: 5’-CCUAAGGUCCGGUUUAUCA-3’; UBE2K#3: 5’-CCAGAAACAUACCCAUUUA-3’ and UBE2K#4: 5’-GCAAAUCAGUACAAACAAA-3’ (a 1:1:1:1 mix of UBE2K siRNAs #1, #2, #3 and #4 was used).

### Immunoblotting, immunoprecipitation and chromatin fractionation

Immunoblotting was performed as previously described ^58^. To prepare cell extracts, cells were lysed in EBC buffer (50 mM Tris, pH 7.5; 150 mM NaCl; 1 mM EDTA; 0.5% NP40; 1 mM DTT) or MDsk lysis buffer (50 mM Tris, pH 8.0; 1 M NaCl; 5 mM EDTA; 1% NP40; 0.1% SDS) supplemented with protease and phosphatase inhibitors on ice for 20 min, and lysates were cleared by centrifugation (16,300g, 20 min). For immunoprecipitation (IP) experiments to analyze protein modification by SUMO and ubiquitin, cells were lysed in denaturing buffer (20 mM Tris, pH 7.5; 50 mM NaCl; 1 mM EDTA; 0.5% NP40; 0.5% SDS, 0.5% sodium deoxycholate; 1 mM DTT) supplemented with protease and phosphatase inhibitors, followed by sonication. For co-IP experiments, cells were lysed in EBC buffer supplemented with protease and phosphatase inhibitors on ice for 20 min. Lysates were then cleared by centrifugation (16,300g, 20 min). Cleared lysates were incubated with GFP-trap or DNMT1-trap agarose (Chromotek) overnight with constant agitation at 4 °C. After washing, immobilized proteins were eluted from the beads by boiling in 2xLaemmli sample buffer for 5 min. For two-step IPs, GFP-tagged DNMT1 was purified on GFP-Trap agarose (Chromotek) under denaturing conditions as described above and washed extensively in denaturing buffer. The beads were equilibrated in EBC buffer and incubated with whole cell lysate prepared in EBC buffer with constant agitation at 4 °C for 4 h. Beads were then washed in EBC buffer and proteins were eluted by boiling in 2xLaemmli sample buffer for 5 min.

For pulldowns of total cellular ubiquitin conjugates using the MultiDsk affinity reagent ^44^, purified Halo-tagged MultiDsk (15 μg per condition) was pre-incubated with HaloLink resin (Promega) for 1 h with rotation at room temperature in binding buffer (100 mM Tris, pH 7.5; 150 mM NaCl; 0.05% IGEPAL). Excess protein was removed by washing with binding buffer supplemented with 1 mg/ml BSA. Cells were lysed in MultiDsk buffer supplemented with protease and phosphatase inhibitors on ice for 15 min, followed by sonication. Lysates were then cleared by centrifugation (16,300g, 20 min). Cleared lysates were incubated with Halo-MultiDsk resin overnight with constant agitation at 4 °C. After multiple washes of the beads, ubiquitylated proteins were eluted from the beads by boiling in 2xLaemmli sample buffer for 5 min.

For chromatin fractionation, cells were first lysed in buffer 1a (10 mM Tris, pH 8.0; 10 mM KCl; 1.5 mM MgCl2; 0.34 M sucrose; 10% glycerol; 0.1% Triton X-100) supplemented with protease and phosphatase inhibitors on ice for 5 min, followed by centrifugation (2000g, 5 min) to recover the soluble proteins. Pellets were then washed extensively once in buffer 1b (buffer 1a supplemented with 500 mM NaCl) and then once in buffer 1a, followed by resuspension in buffer 2 (50 mM Tris, pH 7.5; 150 mM NaCl; 1% NP40; 0.1% SDS; 1 mM MgCl_2_; 125 U/ml benzonase) supplemented with protease and phosphatase inhibitors. Lysates were incubated in a thermomixer (37 °C, 1000 rpm, 15 min), and solubilized chromatin-bound proteins were obtained by centrifugation (16,300g, 10 min).

### Antibodies

Antibodies to human proteins used in this study included: Actin (1:20,000 dilution (MAB1501, Millipore, RRID:AB_2223041)); DNMT1 (1:1,000, described in ^11^); FLAG (1:1,000 (A00187, GenScript, RRID:AB_1720813)); GFP (1:5,000, Abcam (ab6556, RRID:AB_305564)); HA (1:1,000 (11867423001, Roche, RRID:AB_390918)); Histone H3 (1:20,000, Abcam (ab1791, RRID:AB_302613)); PARP1 (1:1,000, Santa Cruz Biotechnology (sc-8007, RRID:AB_628105)); PIAS1 (1:1,000 (ab77231, Abcam, RRID:AB_1524188)); RNF4 (1:5,000, described in ^57^); SUMO1 (1:1,000, Thermo Fisher Scientific (33-2400, RRID:AB_2533109)); SUMO2/3 (1:1,000, Abcam (ab3742, RRID:AB_304041); 1:1,000, Abcam (ab81371, RRID:AB_1658424)); TOPORS (1:250, sheep polyclonal raised against full-length human TOPORS); ubiquitin (1:1,000, Santa Cruz Biotechnology (sc-8017 AC, RRID:AB_2762364); 1:1,000, Millipore (04-263, RRID:AB_612093)); ubiquitin (K48-linked) (1:1,000, Millipore (05-1307, RRID:AB_1587578)); ubiquitin (K63-linked) (1:1,000, (05-1308, RRID:AB_1587580)); UBE2K (1:1,000, Cell Signaling Technology (3847, RRID:AB_2210768)); and USP7 (1:1,000, Bethyl (A300-033A, RRID:AB_203276)).

### Immunofluorescence and high-content image analysis

Cells were pre-extracted on ice in stringent pre-extraction buffer (10 mM Tris–HCl, pH 7.4; 2.5 mM MgCl_2_; 0.5% NP-40; 1 mM PMSF) for 8 min and then in ice-cold PBS for 2 min prior to fixation with 4% formaldehyde for 15 min. If not pre-extracted, cells were subjected to permeabilization with PBS containing 0.2% Triton X-100 for 5 min prior to blocking. Coverslips were blocked in 10% BSA and incubated with primary antibodies for 2 h at room temperature, followed by staining with secondary antibodies and DAPI (Alexa Fluor; Life Technologies) for 1 h at room temperature. Coverslips were mounted in MOWIOL 4-88 (Sigma).

Manual image acquisition was performed with a Zeiss laser scanning confocal microscope LSM880, using the Plan-Apochromat 40× 1.30 Oil DIC M27 objective and ZEN Software (Zeiss). Coverslips were prepared as described above, except that they were mounted with Vectashield Mounting medium (Vector Laboratories) and sealed with nail polish. Raw images were exported as TIFF files and processed using Adobe Photoshop. If adjustments in image contrast and brightness were applied, identical settings were used on all images of a given experiment. For automated image acquisition, images were acquired with an Olympus IXplore ScanR system equipped with Olympus IX-83 wide-field microscope, Yokogawa CSU-W1Confocal spinning disk unit, four 50/100mW laser diodes, UPLSXAPO20× NA 0.80 WD 0.60 mm or UPLSXAPO40×2 NA 0.95 WD 0.18 mm, and Hamamatsu Orca Flash 4 sCMOS camera. Automated and unbiased image analysis was carried out with the ScanR analysis software. Data were exported and processed using Spotfire (TIBCO Software Inc.).

### Live-cell imaging

U2OS cells stably expressing Fucci (fluorescent, ubiquitination-based cell cycle indicator) reporter constructs (mKO2-hCdt1(30-120) and mAG-hGem(1-110)) ^46^ were transfected with siRNAs and seeded onto an Ibidi dish (Ibidi) 16 h before acquisition. Medium were changed to Leibovitz’s L15 medium (Life Technologies) supplemented with 10% FBS prior to filming. Live-cell imaging was performed on a Deltavision Elite system using a 40× oil objective (GE Healthcare). Images were acquired every 10 or 15 min for 48 h. Three z-stacks of 5 μm were imaged. SoftWork software (GEHealthcare) were used for data analysis.

### Proliferation and survival assays

Relative proliferation was measured using an Incucyte (Sartorius) instrument. Cells (6×10^4^) were seeded into a 24- or 96-well plate 24 h after siRNA treatment and imaged every 2 h. Mean confluency was determined from four (**Figure 5E**) or nine (**Figure S5F,G**) images. For survival assays, approx. 400 cells were plated per 60-mm plate and treated with 5-AzadC for 24 h or formaldehyde for 30 min. Cells were then washed with PBS twice and fresh medium was added. Colonies were fixed and stained after approximately 2 weeks with cell staining solution (0.5% w/v crystal violet, 25% v/v methanol). The number of colonies were quantified using a GelCount (Oxford Optronix) colony counter.

### Purification of recombinant TOPORS proteins

HEK293-6E suspension cells were cultured in FreeStyle F17 expression medium (Gibco) supplemented with 4 mM L-glutamine (Gibco), 1% FBS, 0,1% Pluronic F-68 (Gibco) and 50 μg/ml G418 (Invivogen) in a 37 °C incubator on a shaker rotating at 140 rpm. HEK293-6E cells were transfected with Strep-HA-TOPORS WT, *SIM or *RING expression plasmids using PEI transfection reagent (Polyscience Inc). After 24 h, cells were harvested and snap-frozen in liquid nitrogen. The cell pellet was resuspended in lysis buffer (100 mM Tris, pH 8.0; 350 mM NaCl; 0,05% NP-40 and protease inhibitors), benzonase-treated and sonicated. The lysate was then cleared by centrifugation at 25,000g at 4 °C for 30 min. The cleared lysate was incubated with Strep-Tactin Superflow resin (IBA) and incubated at 4 °C for 2 h. After the beads were washed in a gravity column using wash buffer (100 mM Tris, pH 8.0; 600 mM NaCl; 0,05% NP-40), TOPORS protein was eluted using elution buffer (100 mM Tris, pH 8.0; 350 mM NaCl; 2,5 mM Desthiobiotin). The elute fractions were run on a 4-12% NuPAGE Bis-Tris protein gel (Invitrogen) and stained with Instant Blue Coomassie Protein Stain (Expedeon). Fractions were concentrated on Microcon-30 kDa Centrifugal Filters (Millipore), snap-frozen in liquid nitrogen and stored at −70°C. For *in vitro* ubiquitylation and SUMOylation reactions, 0.35 μg Strep-HA-TOPORS prep was used, for Strep-Tactin pull-downs 0.7 μg was used, and for STUbL assays 2.8 μg was used.

### In vitro ubiquitylation, SUMOylation and deubiquitylation reactions

All *in vitro* ubiquitylation and SUMOylation reactions were carried out in reaction buffer containing 50 mM Tris pH 7.5; 150 mM NaCl; 0.1% NP-40; 5 mM MgCl_2_ and 0.5 mM TCEP. Unless otherwise stated, the following final concentrations were used: FLAG Ubiquitin (R&D Systems, 20 μM), ATP (Sigma-Aldrich, 3 mM), UBA1 (100 nM), UBE2D1 (0.5 μM), UBE2K (R&D Systems, 0.28 μM), SAE1-UBA2 (R&D Systems, 100 nM), UBE2I (R&D Systems, 0.5 μM), SUMO2 (R&D Systems, 2 μM), poly-SUMO2_2-8_ chains (R&D Systems, 0.3 μg per reaction), RNF4 (47 nM), GST-PIAS1 (Enzo Life Science), 4xSUMO2 and Ub-4xSUMO2 (1.8 μM) ^34^. HA-SUMO1 Vinyl Sulfone (VS) and HA-Ubiquitin VS (R&D Systems) were added to all reactions containing purified Strep-HA-TOPORS proteins in order to inhibit deubiquitinase and SUMO protease activities co-purifying with Strep-HA-TOPORS.

For *in vitro* deubiquitylation reactions with recombinant USP7, GFP-TOPORS expressed in U2OS cells was immunoprecipitated using GFP-trap agarose under denaturing condition as above, followed by extensive washing. Beads containing bound GFP-TOPORS were equilibrated in deubiquitylation buffer (50 mM Tris, pH 7.5; 150 mM NaCl; 5 mM DTT) and incubated with 0.5 µg recombinant USP7 protein (Ubiquigent) with shaking at 30 °C for 30 min. Beads were washed, eluted by boiling in 2xLaemmli sample buffer for 5 min and analyzed by immunoblotting.

### Flow cytometry

Cells were collected by trypsinization, fixed in 4% paraformaldehyde in PBS for 15 min and permeabilized in 0.2% Triton-X, 2% FBS in PBS for 20 min at room temperature. Permeabilized cells were washed in FACS buffer (PBS+10% FBS) and incubated with sheep polyclonal anti-DNMT1 antibody ^14^ diluted in FACS buffer for 90 min at room temperature. Following two washes in FACS buffer, cells were incubated with anti-sheep AlexaFluor 488 (Invitrogen) diluted in FACS buffer for 1 h at room temperature. For EdU co-staining, washed cells were subsequently stained using the Click-iT Plus EdU Alexa Fluor 647 Kit (Invitrogen) according to the manufacturer’s instructions. Following two additional washes in FACS buffer, cells were resuspended in FACS buffer containing 1 µg/ml DAPI (Thermo Fisher), strained to a single cell solution (40 µm filter), and analyzed using a BD LSRFortessa flow cytometer (BD Biosciences). Analysis was performed and plots were generated in FlowJo software.

### Mutagenesis of HAP1 cells

Gene-trap mutagenesis of HAP1 cell lines were carried out as described ^59^. Briefly, a BFP-containing variant of gene-trap retrovirus was generated in low passage HEK293T cells by co-transfection of gene-trap vector, retroviral packaging plasmids Gag-pol and VsVg, and pAdvantage (Promega). Media containing retrovirus was collected 48 h and 72 h after transfection, concentrated using centrifugation filters (Amicon) and the pooled, concentrated retrovirus was used to transduce 40×10^6^ HAP1 cells supplemented with protamine sulfate (8 µg/mL) for 24 h. After recovery, mutagenized HAP1 cells were expanded and used for genetic screens.

### FACS-based screens for DNMT1 abundance

To identify regulators of 5-AzadC-induced DNMT1 processing, mutagenized HAP1 WT cells were expanded to 3×10^9^ cells per screen. Two screens were performed: a mock DNMT1 screen where cells were labelled for 2 h with 10 µM EdU and a perturbation DNMT1 screen with co-treatment of 10 µM 5-AzadC and EdU. Treated cells were then trypsinized, fixed, permeabilized and stained for DNMT1 and EdU as described in the flow cytometry section above. Cells were sorted on a BD FACSAria Fusion cell sorter gating for haploid EdU-incorporating cells and the cells with the 5% highest and lowest DNMT1 signal, respectively, were collected. Genomic DNA was extracted from sorted cells (1,3×10^7^ cells per channel) using the QIAamp DNA Mini kit (Qiagen). Gene-trap insertion site recovery and sequencing libraries were generated as previously described ^60^.

Amplified libraries were sequenced on a HiSeq2500 (Illumina) with a read length of 65 bp (single-end read). Sequencing reads from each sample (low and high) were aligned to the human genome (hg38) allowing one mismatch and assigned to non-overlapping protein-coding gene regions (Refseq). The number of unique gene-trap insertions in the sense direction of each gene was normalized to the total number of sense insertions of each sorted population. The mutational index (MI) was calculated for each gene by comparing the number of unique, normalized sense integrations of the high population to that of the low population using a two-sided Fisher’s exact test (FDR-corrected p<0.05). Every gene was then plotted in Fishtail scatterplots comparing the combined number of unique insertions identified in the two populations of a given gene (log10, x-axis) to its mutational index (log2, y-axis). To identify genes selectively affecting DNMT1 abundance in response to 5-AzadC, but not in the mock screen, a comparative filtering of genes was performed. Significant positive and negative regulators that scored as such in both screens (except for the antigen target DNMT1) were removed from the 5-AzadC screen to highlight DNMT1 DPC-specific regulators and generate **Figure 1B**. Significant regulators for each individual screen can be found in **Table S1**. Fishtail scatterplots were generated using GraphPad Prism software.

### Fitness-based screens to identify synthetic lethal interactions

Haploid genetic fitness screens were carried out as described in ^45^. Briefly, mutagenized HAP1 cell lines (minimum coverage of 2,5×10^8^ cells per screen) were passaged for 10 days, trypsinized and fixed using Fixation buffer I (BD Biosciences) for 10 min at 37 °C. For RNF4-dTAG cells, cells were passaged in the presence of 0.25 µM dTAG13 to induce loss of RNF4. Cells were then permeabilized with Perm buffer III (BD Biosciences) and stained in FACS buffer containing 2.5 µg/ml DAPI (Thermo Fisher). After washing in FACS buffer, cells were strained to a single cell solution (40 µm filter), and a minimum of 3×10^7^ haploid cells (based on DAPI content) isolated using a BD FACSAria Fusion cell sorter. Isolation of genomic DNA, library preparation and insertion site mapping were done as described for the FACS-based screens above. Analysis of gene-trap insertion orientation bias (sense-ratio) was performed as described ^45^, and the analysis pipeline can be found on GitHub (https://github.com/BrummelkampResearch/phenosaurus). For each replicate experiment a two-sided binomial test was calculated which gives a p-value for each gene. These p-values were then corrected for False Discovery Rate (FDR) using the Benjamin-Hochberg Procedure and the least significant p-value among the individual replicates of a genotype was used to determine if a gene is considered significant. Every replicate corresponds to an independent clonal cell line of the respective genotype. Four independent cultured wild-type control datasets published in ^45^ were used as control (available at SRA SRP058962, accession numbers SRX1045464, SRX1045465, SRX1045466, SRX1045467). To identify genes that affect cell viability selectively in TOPORS- and RNF4-deficient cell lines, the number of disruptive sense integrations and non-disruptive antisense integrations for each gene was compared to that in the four control datasets using a two-sided Fisher’s exact test. Genes with a significant orientation bias in screen replicates of TOPORS- and RNF4-deficient cells, respectively, in addition to a significantly altered sense-ratio (p < 0.05, odds ratio < 0.8) in relation to the control datasets were considered as hits.

### Generation of HAP1 cell lines

To generate clonal TOPORS knockout cell lines, parental HAP1 cells were co-transfected with a blasticidin resistance cassette and the CRISPR-Cas9 vector pX330 containing sgRNA sequences targeting *TOPORS* (sgTOPORS#1 (5’-AACAGTACTCCACTATCCGG-3’) and sgTOPORS#2 (5’-GGTAGCGAAATCGTCGATCA-3’)). Cells were then briefly selected (48 h) during clonal outgrowth and gene status monitored using Sanger sequencing of genomic DNA and immunoblot analysis. C-terminal degron tagging of endogenous RNF4 was carried out by a generic CRISPR-Cas9 strategy as described ^61^, but using a modified pTIA donor vector containing HA-tagged FKBP12(F36V) (dTAG) and a P2A sequence followed by a blasticidin cassette for integration selection (pTIA dTAG-HA P2A Blast). To generate clonal cell lines, parental HAP1 cells were co-transfected (1:1) with pTIA dTAG donor vector and a CRISPR-Cas9 vector containing a sgRNA targeting the last exon (exon 8) of RNF4 (pX330 sgRNF4 (5’-TACTTCATATATAAATGGGG-3’)) and subjected to blasticidin selection during clonal outgrowth. HAP1 RNF4-dTAG clones were validated by Sanger sequencing of the genomic locus and immunoblot analysis. The two RNF4-dTAG clones used contained an in-frame insertion of the dTAG donor vector at the C-terminus of RNF4, after amino acid His186 (hg38 genome coordinate chr4:2,513,804).

### Mass spectrometry analysis of DNMT1 pulldowns

U2OS cells transfected with siRNAs for 72 h were synchronized by single-round thymidine treatment. Cells were released from synchronization and treated or not with 5-AzadC for 30 min in early S phase. Cells were then collected and lysed in denaturing buffer (20 mM Tris, pH 7.5; 50 mM NaCl; 1 mM EDTA; 0.5% NP40; 0.5% SDS, 0.5% sodium deoxycholate; 1 mM DTT) and subjected to IP using DNMT1-trap beads, as described above. After overnight incubation, beads were washed extensively in denaturing buffer. To make samples MS-compatible, beads were washed with minimal washing buffer (50 mM Tris, pH 7.5; 150 mM NaCl) for 15 min and transferred to Protein LoBind tubes (Eppendorf). The washing procedure was repeated twice and beads were again transferred to new Protein LoBind tubes. Beads were then washed one final time with minimal washing buffer (at pH 8.0) for 15 min.

Digestion of proteins was performed on-beads, using 250 ng (5 ng/µl) trypsin, with 1 h pre-incubation on ice and overnight incubation with shaking at 30 °C. Digests were cleared by centrifugation through 0.45 µm spin filters, after which peptides were reduced and alkylated using chloroacetamide and tris(2-carboxyethyl)phosphine at final concentrations of 5 mM. Peptides were purified on StageTips at high pH, with C18 StageTips prepared in-house, by layering four plugs of C18 material (Sigma-Aldrich, Empore SPE Disks, C18, 47 mm) per StageTip. Activation of StageTips was performed with 100 μL 100% methanol, followed by equilibration using 100 μL 80% acetonitrile (ACN) in 200 mM ammonium hydroxide, and two washes with 100 μL 50 mM ammonium hydroxide. Samples were basified to pH >10 by addition of one fifth volume of 200 mM ammonium hydroxide, after which they were loaded on StageTips. Subsequently, StageTips were washed twice using 100 μL 50 mM ammonium hydroxide, after which peptides were eluted using 80 µL 25% ACN in 50 mM ammonium hydroxide. All fractions were dried to completion using a SpeedVac at 60 °C. Dried peptides were dissolved in 11 μL 0.1% formic acid and stored at −20 °C until analysis using mass spectrometry (MS).

All samples were analyzed on an EASY-nLC 1200 system (Thermo) coupled to an Orbitrap Exploris 480 mass spectrometer (Thermo). Samples were analyzed on 20 cm long analytical columns, with an internal diameter of 75 μm, and packed in-house using ReproSil-Pur 120 C18-AQ 1.9 µm beads (Dr. Maisch). The analytical column was heated to 40 °C, and elution of peptides from the column was achieved by application of gradients with stationary phase Buffer A (0.1% formic acid (FA)) and increasing amounts of mobile phase Buffer B (80% ACN in 0.1% FA). The primary analytical gradient ranged from 5 %B to 32 %B over 60 min, followed by a tail-end increase to 42 %B over 5 min to ensure full peptide elution, followed by a washing block of 15 min. Ionization was achieved using a NanoSpray Flex NG ion source (Thermo), with spray voltage set at 2 kV, ion transfer tube temperature to 275 °C, and RF funnel level to 40%. Full scan range was set to 300-1,300 m/z, MS1 resolution to 120,000, MS1 AGC target to “200” (2,000,000 charges), and MS1 maximum injection time to “Auto”. Precursors with charges 2-6 were selected for fragmentation using an isolation width of 1.3 m/z and fragmented using higher-energy collision disassociation (HCD) with normalized collision energy of 25. Monoisotopic Precursor Selection (MIPS) was enabled in “Peptide” mode. Precursors were prevented from being repeatedly sequenced by setting expected peak width to 40 s, and setting dynamic exclusion duration to 80 s, with an exclusion mass tolerance of 15 ppm, exclusion of isotopes, and exclusion of alternate charge states for the same precursor. MS/MS resolution was set to 45,000, MS/MS AGC target to “200” (200,000 charges), MS/MS intensity threshold to 230,000, MS/MS maximum injection time to “Auto”, and TopN to 9.

### Mass spectrometry data analysis (DNMT1 pulldowns)

All RAW files were analyzed using MaxQuant software (version 1.5.3.30) ^62,63^. Default MaxQuant settings were used, with exceptions outlined below. For generation of the theoretical spectral library, the HUMAN.FASTA database was extracted from UniProt on 3 September, 2021. Protein N-terminal acetylation (default), methionine oxidation (default), and lysine ubiquitylation (i.e. GlyGly), were included as potential variable modifications, with a maximum allowance of 3 variable modifications per peptide. First search mass tolerance was set to 10 ppm, and maximum charge state of considered precursors to 6. Label-free quantification (LFQ) was enabled, and “Fast LFQ” was disabled. Second peptide search was enabled (default) and matching between runs was enabled with a match time window of 1 min and an alignment time window of 30 min. Note that the alignment time window was increased (from default 20 min) because during one run (“20221116_EXPL7_LC6_IAH_collab_JL_D-IP_siD_AZA_01.raw”) contact closure initiated early, resulting in the mass spectrometer starting the recording during sample loading, causing all peaks in the gradient to be shifted by ∼15 min. Minimum Score and minimum Delta Score for modified peptides were increased to 80 and 40, respectively, and site decoy fraction was set to 0.02. Data were filtered by posterior error probability to achieve a false discovery rate of <1% (default), at both the peptide-spectrum match and the protein assignment levels.

### MS data statistics (DNMT1 pulldowns)

Statistical handling of MS data was performed using Perseus software (versions 1.5.5.3 and 1.6.14.0) ^64^, and visualized using GraphPad Prism (version 9.5.1). MS data was filtered by removing reverse-database hits and potential contaminants. Following 2log transformation, only proteins and peptides observed at n=4/4 in at least one experimental condition were considered. In case of protein-level quantification, missing values were globally imputed (down shift 1.8, width 0.3).

### Proteomic analysis of PML body components

The generation of U2OS PML-KO/YFP-PML cells and purification of PML bodies has been described previously ^38^. The purification procedure here was followed exactly as detailed, except that cells were grown in SILAC medium for at least 8 doublings prior to expansion from a single 75 cm^2^ flask to five confluent 15 cm dishes. Cells were treated with 1 µM arsenic trioxide for 2 h (Heavy condition) or left untreated (Light condition). Purified PML body samples from each SILAC condition were mixed in 1:1 ratio by protein mass, then fractionated by SDS-PAGE and the lane cut into 7 slices. Tryptic peptides were extracted and approximately 15-30% of the total peptide yield for each slice was analyzed by LC-MS/MS using a 150-min elution gradient and a top 10 data-dependent method on a Q Exactive mass spectrometer broadly as described previously ^38^.

The remaining tryptic peptides were further digested with GluC and approximately half of the yield was analyzed by LC-MS/MS using a 180-min elution gradient and similar top 10 data-dependent mass spectrometer settings. MS data were processed in MaxQuant version 1.6.1.0 ^65,66^ using a UniProt human database containing 73,920 proteins downloaded in 2019 and the YFP-PML sequence. Oxidized methionine and acetylated protein N-termini were selected as variable modifications and carbamidomethyl-C was a fixed modification. Digestion assumed Trypsin/P (maximum missed cleavages of 2) or Trypsin/P plus GluC (maximum missed cleavages of 5) depending on the peptide preparations analyzed. Match between runs and advanced ratio estimation were selected. Using 1% FDR filtering against a reverse decoy database at peptide and protein levels, data for 1,772 protein groups were returned which was reduced to 1,025 by removal of decoy proteins, proteins only identified by modified peptides, proteins from the contaminants database, proteins without a reported H/L ratio and proteins with fewer than three unique peptides.

## Acknowledgements

We thank Abdelghani Mazouzi and members of the Mailand laboratory for helpful discussions, and Guizela H. Prince for bioinformatic support. This work was supported by grants from Novo Nordisk Foundation (grants no. NNF14CC0001 and NNF18OC0030752), Lundbeck Foundation (grant no. R223-2016-281), Independent Research Fund Denmark (grant no. 0134-00048B) and Danish National Research Foundation (grants no. DNRF-115 and DNRF-166). R.T.H. was supported by an Investigator Award from Wellcome Trust (217196/Z/19/Z) and, with M.H.T., a Programme grant from Cancer Research UK (C434/A21747). J.C.Y.L. was supported by the Croucher Foundation. P.H. was supported by Novo Nordisk Foundation.

## Author Contributions

Conceptualization: J.C.Y.L., P.H. and N.M.; Methodology: J.C.Y.L., S.H., I.A.H., M.H.T., G.M., R.T.H., M.L.N., T.B., P.H. and N.M.; Investigation: J.C.Y.L., L.A., S.H., Z.G., I.A.H., C.J., L.M., M.H.T. and P.H.; Writing – Original Draft: N.M.; Writing – Review and Editing: All authors; Supervision, R.T.H., M.L.N., T.B. and N.M.; Project Administration: P.H. and N.M.; Funding Acquisition: R.T.H., P.H. and N.M.

## Conflict of Interest

The authors declare no competing interests.

## Supplemental figure legends

**Figure S1 (related to Figure 1).**
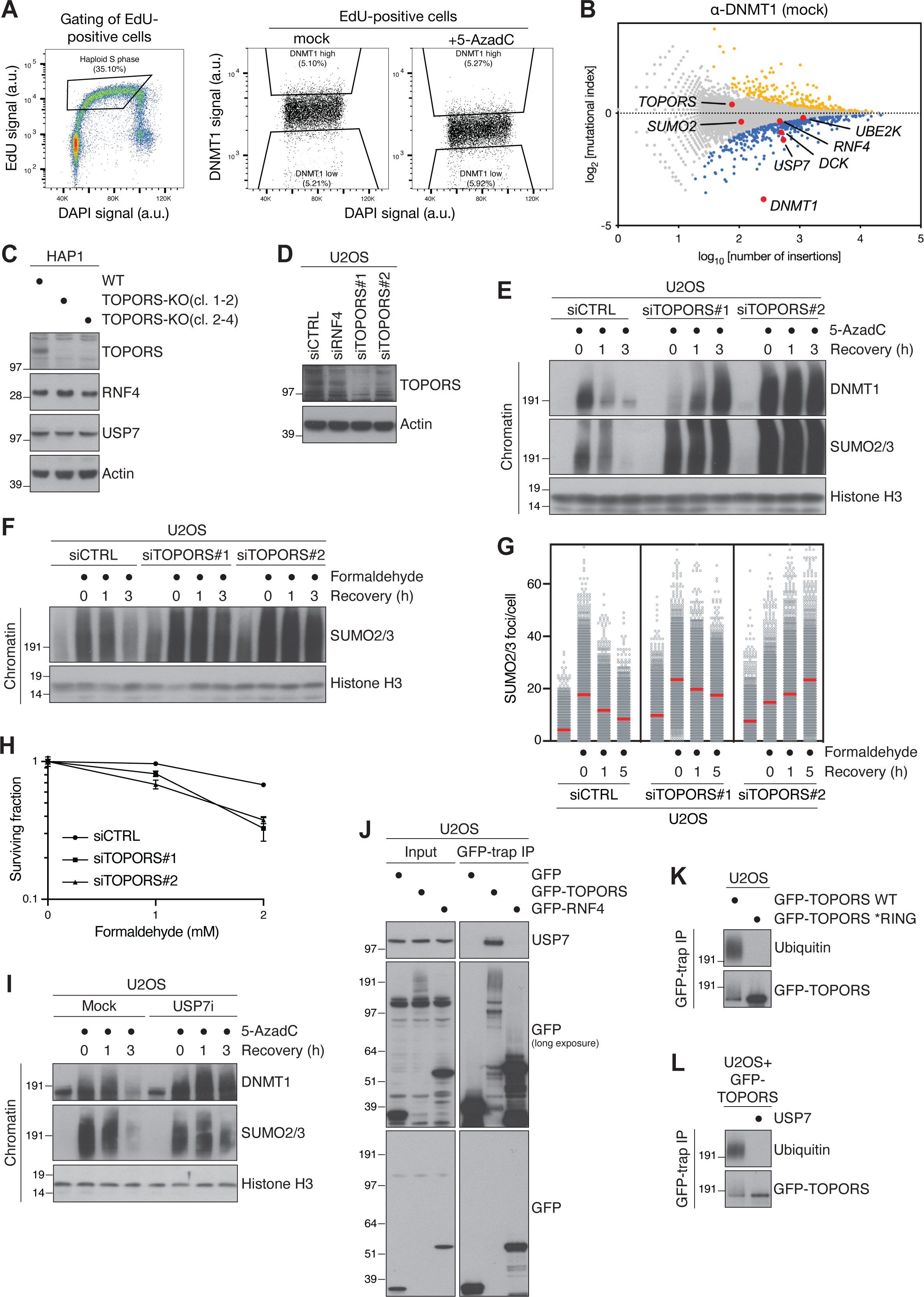
TOPORS is required for SUMO-dependent DPC resolution and is stabilized by USP7. **A.** Flow cytometry plots of the gating strategy used for FACS-based DNMT1 haploid genetic screens in Figure 1B and **Figure S1B**. **B.** Mock screen for DNMT1 abundance in EdU-positive cells, presented as a fishtail plot in which genes are plotted according to their mutational index (y-axis) and the total number of gene-trap insertions in the gene (x-axis). Positive and negative regulators are labeled in blue and yellow, respectively (two-sided Fisher’s exact test, FDR corrected p≤0.05; non-significant genes are shown in grey). **C.** Immunoblot analysis of HAP1 WT and TOPORS-KO cells. **D.** Immunoblot analysis of U2OS cells transfected with non-targeting control (CTRL), RNF4, or TOPORS siRNAs. **E.** U2OS cells transfected with indicated siRNAs were released from thymidine synchronization, left untreated or exposed to 5-AzadC for 30 min and collected at the indicated times. Chromatin-enriched fractions were immunoblotted with indicated antibodies. **F.** Immunoblot analysis of chromatin-enriched fractions of U2OS cells transfected with indicated siRNAs, left untreated or exposed to formaldehyde for 1 h and collected at the indicated times. **G.** Cells treated as in (F) were pre-extracted and stained with SUMO2/3 antibody. SUMO2/3 foci count per cell was analyzed by QIBC (red bars, mean; >4,400 cells analyzed per condition). Data are representative of 3 independent experiments. **H.** Clonogenic survival of U2OS cells transfected with indicated siRNAs and subjected to indicated doses of formaldehyde for 30 min before replating (mean±SEM; *n*=3 independent experiments). **I.** As in (E), except that USP7i was administered together with 5-AzadC where indicated. **J.** U2OS cells transfected with indicated GFP expression plasmids were subjected to GFP IP and immunoblotted with indicated antibodies. **K.** U2OS cells transfected with indicated GFP-TOPORS expression constructs and subjected to GFP IP under denaturing conditions and immunoblotted with indicated antibodies. Mutations in the TOPORS RING domain (*RING) that inactivate its E3 ubiquitin ligase activity (see Figure 2E) abolish TOPORS ubiquitylation, suggesting it is modified by auto-ubiquitylation. **L.** U2OS cells transfected with GFP-TOPORS WT expression plasmid were subjected to GFP IP under denaturing conditions. Immobilized proteins were subjected to stringent washing, incubated with or without recombinant USP7 protein at 30 °C for 30 min and immunoblotted with ubiquitin and GFP antibodies.

**Figure S2 (related to Figure 2).**
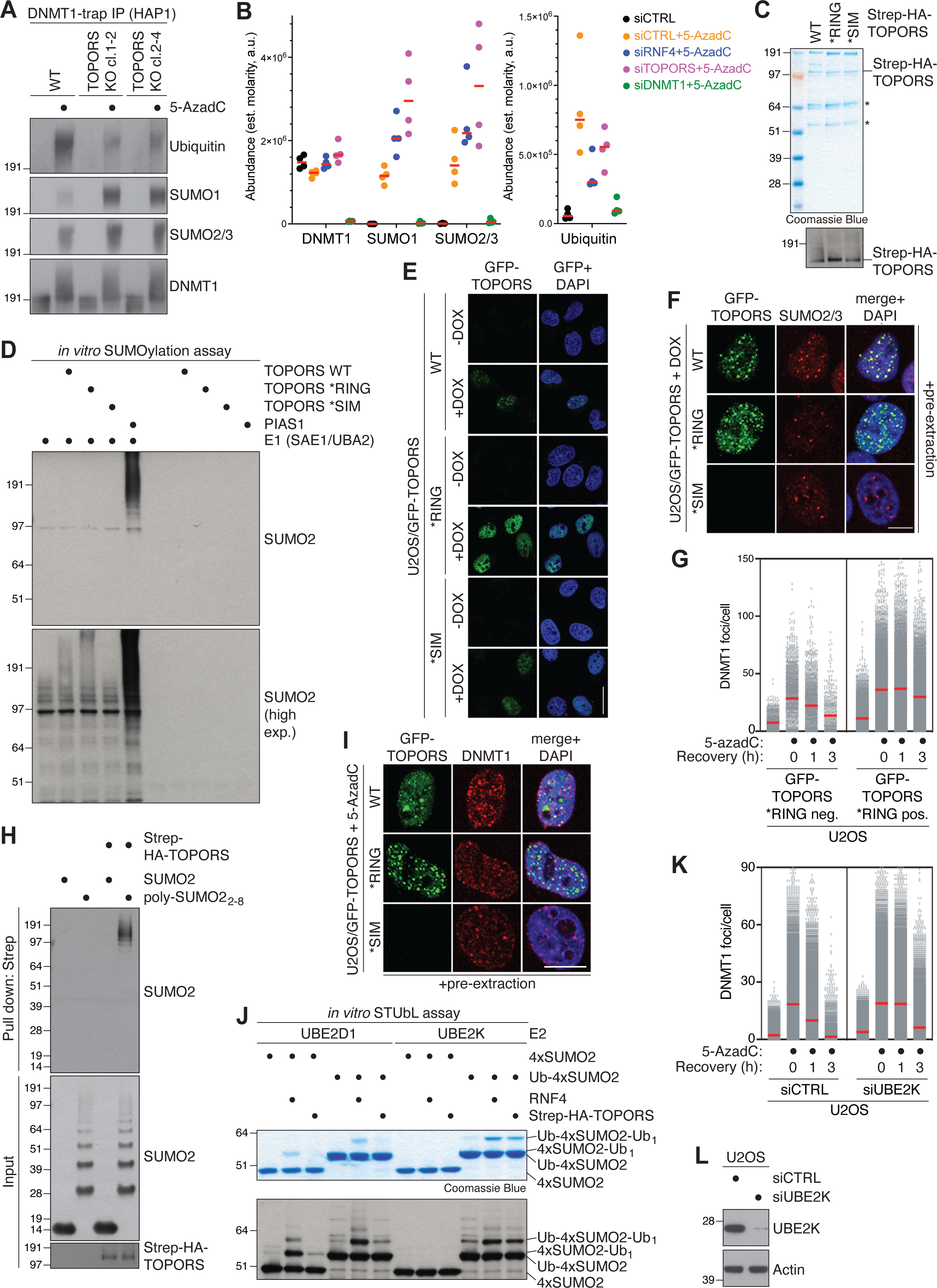
TOPORS functions as a SUMO-targeted ubiquitin ligase in DPC repair in conjunction with UBE2K. **A.** Immunoblot analysis of HAP1 WT and TOPORS-KO cells transfected with non-targeting control (CTRL) or TOPORS siRNAs for 72 h, treated or not with 5-AzadC for 30 min, collected and subjected to DNMT1 IP under denaturing conditions. **B.** Mass spectrometry (MS)-based quantification of DNMT1, SUMO1, SUMO2/3, and ubiquitin in DNMT1 IPs. U2OS cells transfected with indicated siRNAs were subjected to DNMT1 IP under denaturing conditions to isolate DNMT1 but not associated proteins. Samples were digested with trypsin, after which peptides were purified and identified by MS (*n*=4 independent experiments). Molarity was approximated by dividing each protein’s intensity-based abundance by its molecular weight. **C.** Strep-HA-TOPORS proteins purified from HEK293-6E cells were subjected to SDS-PAGE and stained with Coomassie Blue (top) or immunoblotted with HA antibody. Asterisks indicate Strep-HA-TOPORS degradation products. **D.** Immunoblot analysis of *in vitro* SUMOylation reactions containing recombinant Strep-HA-TOPORS proteins (**Figure S2C**) that were pre-incubated with SUMO2-VS in reaction buffer for 10 min, supplemented with E1 (SAE1-UBA2) and E2 (UBE2I) enzymes, SUMO2 and ATP and incubated at 30 °C for 1 h. **E.** Representative images of stable U2OS/GFP-TOPORS cell lines induced or not to express GFP-TOPORS proteins with Doxycycline (DOX). Cells were not pre-extracted prior to fixation and microscopy. Scale bar, 10 μm. **F.** Representative images of DOX-treated U2OS/GFP-TOPORS cell lines immunostained with SUMO2/3 antibody after pre-extraction and fixation. Scale bar, 10 μm. **G.** U2OS cells stably expressing GFP-TOPORS *RING were released from single thymidine synchronization in early S phase, and pulse-treated with 5-AzadC for 30 min. Cells were collected at indicated time points, pre-extracted and immunostained with DNMT1 antibody. DNMT1 foci formation in GFP-positive and -negative cells was analyzed by QIBC (red bars, mean; >479 cells analyzed per condition). Data are representative of three independent experiments. **H.** Immunoblot analysis of recombinant Strep-HA-TOPORS proteins that were incubated with free SUMO2 or poly-SUMO2 chains and subjected to Strep-Tactin pulldown. **I.** Representative images of DOX-treated U2OS/GFP-TOPORS cell lines immunostained with DNMT1 antibody after pre-extraction and fixation. Scale bar, 10 μm. **J.** Coomassie staining (top) and immunoblot analysis using SUMO2 antibody (bottom) of *in vitro* STUbL reactions containing recombinant RNF4 or Strep-HA-TOPORS proteins that were pre-incubated with Ub-VS and SUMO2-VS in reaction buffer for 10 min, supplemented with E1 (UBA1) and E2 (UBE2D1 or UBE2K as indicated) enzymes, FLAG-ubiquitin, 4xSUMO2 STUbL substrate and ATP and incubated at 37 °C for 2 h. **K.** U2OS cells transfected with indicated siRNAs were released from single thymidine synchronization in early S phase and pulse-treated with 5-AzadC for 30 min. Cells were collected at indicated time points, pre-extracted and immunostained with DNMT1 antibody. DNMT1 foci formation was analyzed by QIBC (red bars, mean; >6,400 cells analyzed per condition). Data are representative of three independent experiments. **L.** Immunoblot analysis of U2OS cells transfected with non-targeting control (CTRL) or UBE2K siRNAs.

**Figure S3 (related to Figure 3).**
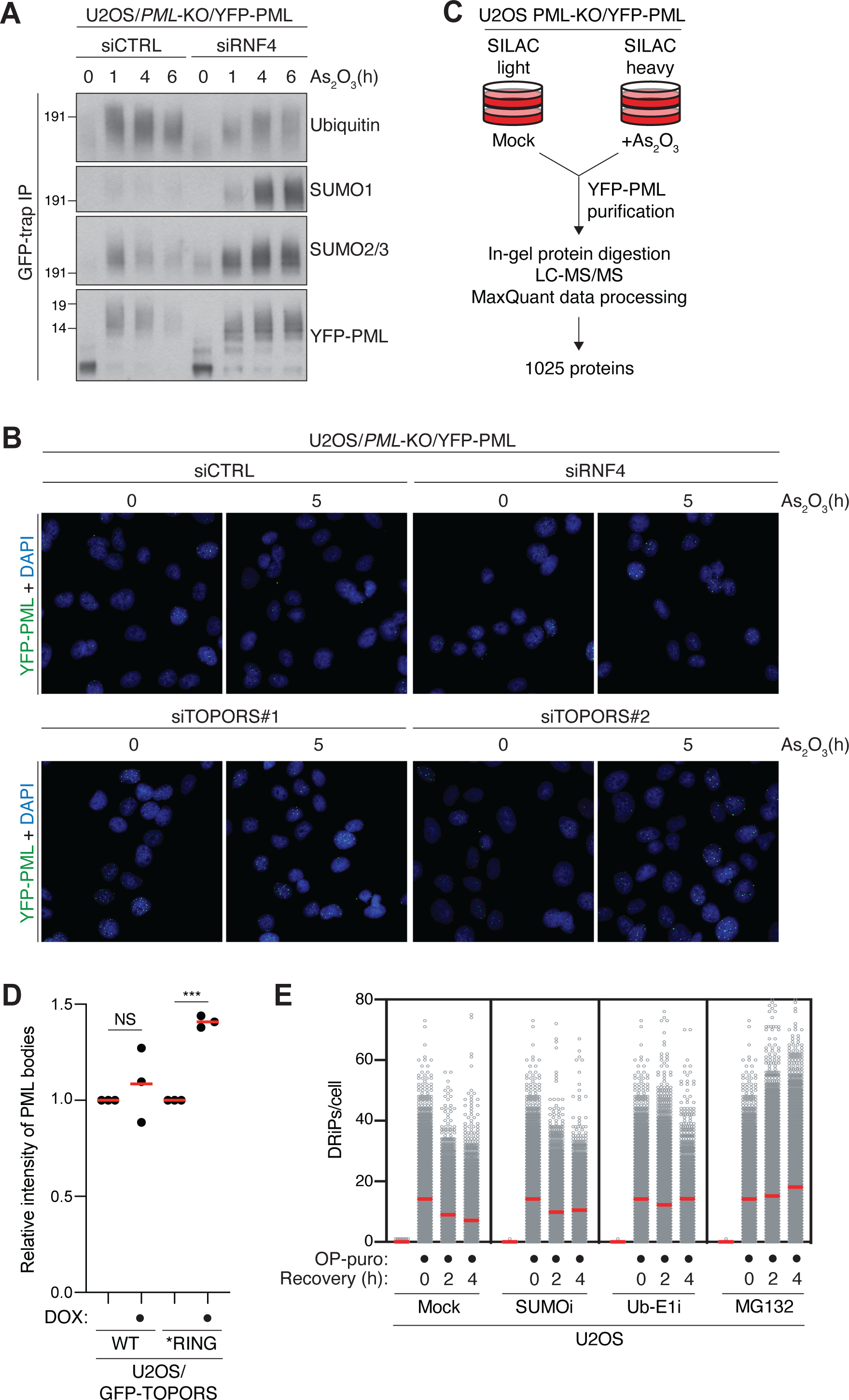
STUbL-dependent turnover of PML bodies and DRiPs. **A.** U2OS PML-KO/YFP-PML cells transfected with indicated siRNAs were treated with arsenic and collected at indicated time points. Cells were then subjected to GFP IP under denaturing conditions and immunoblotted with indicated antibodies. **B.** Representative images of cells in Figure 3B. **C.** Schematic overview of the proteomics experiment conducted for the data described in Figure 3E. **D.** U2OS cells stably expressing GFP-TOPORS WT or *RING were treated or not with Doxycycline (DOX) for 36 h. Cells were pre-extracted and immunostained with PML antibody. Intensity of PML bodies in GFP-positive and -negative cells was analyzed by QIBC and normalized to untreated GFP-negative cells (red bars, mean; *n*=3; ***p<0.001, NS: not significant, one-tailed paired t-test). **E.** U2OS cells were treated with MG132 and O-Propargyl-puromycin (OP-puro) for 4 h, washed, released into fresh media or media containing SUMOi, Ub-E1i or MG132, and collected at the indicated times. OP-puro was click-conjugated to fluorescent azide and analyzed by QIBC (red bars, mean; >2,300 cells analyzed per condition). Data are representative of three independent experiments.

**Figure S4 (related to Figure 4).**
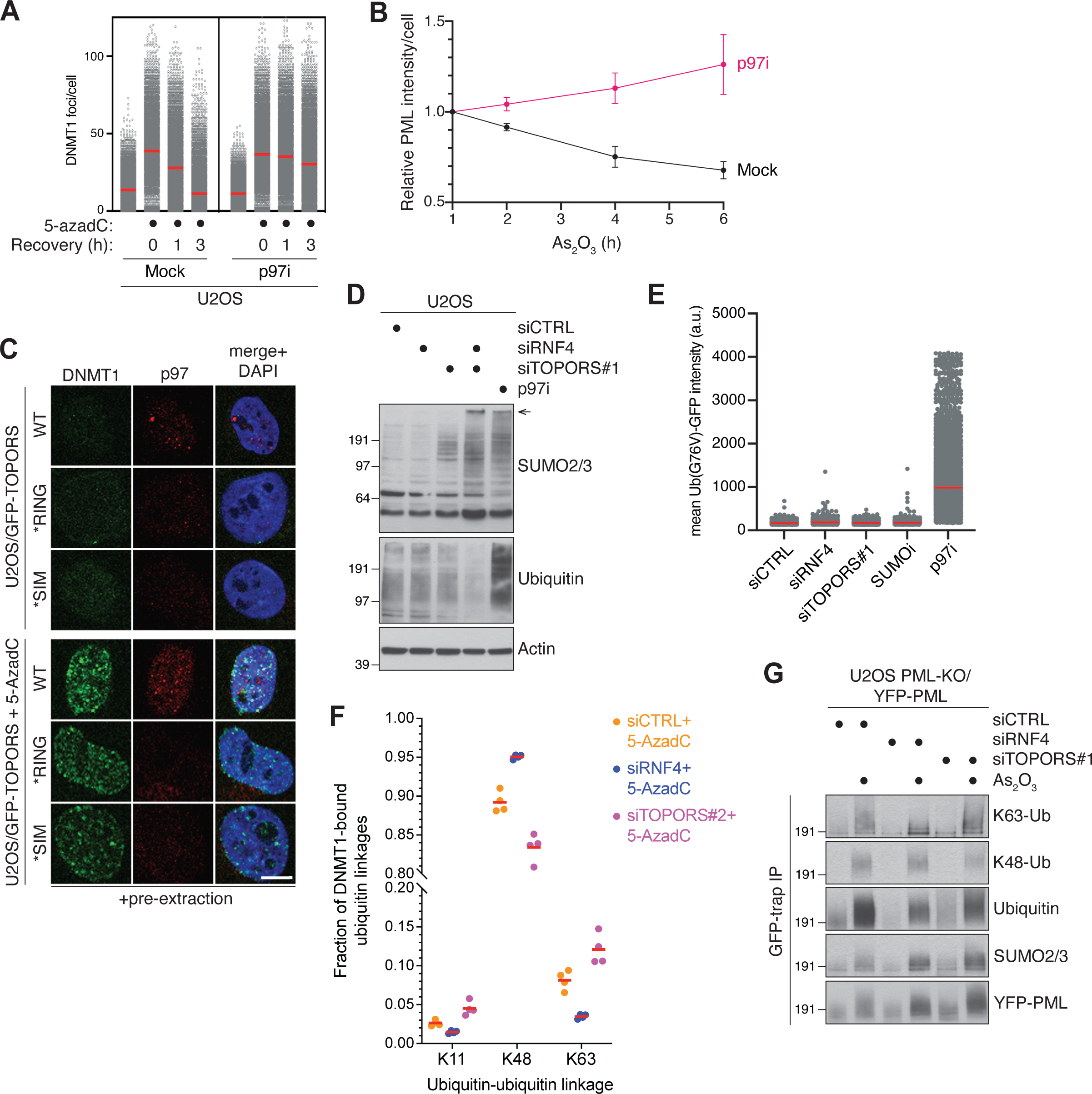
RNF4 and TOPORS promote p97-dependent processing of SUMOylated proteins. **A.** U2OS cells released from single thymidine synchronization in early S phase were treated or not with p97 inhibitor (p97i), pulse-labeled with 5-AzadC for 30 min and collected at the indicated times. Cells were pre-extracted and immunostained with DNMT1 antibody, and DNMT1 foci formation was analyzed by QIBC (red bars, mean; >8,100 cells analyzed per condition). Data are representative of three independent experiments. **B.** U2OS PML-KO/YFP-PML cells were exposed to arsenic in the absence or presence of USP7i. Cells were collected at indicated time points, pre-extracted and YFP-PML intensity was analyzed by QIBC (mean±SEM; *n*=3 independent experiments). **C.** Representative images of DOX-treated U2OS/GFP-TOPORS cell lines that were treated or not with 5-AzadC and co-immunostained with DNMT1 and p97 antibodies after pre-extraction and fixation. Scale bar, 10 μm. **D.** Immunoblot analysis of U2OS cells transfected with indicated siRNAs for 40 h or treated with p97i for 10 h. **E.** U2OS stably expressing Ub(G76V)-GFP reporter transfected with indicated siRNAs were treated or not with p97i for 4 h, and nuclear Ub(G76V)-GFP signal intensity was analyzed by QIBC (red bars, mean; >12,000 cells analyzed per condition). Data are representative of two independent experiments. **F.** Mass spectrometry (MS)-based analysis of ubiquitin-ubiquitin linkages on DNMT1. U2OS cells transfected with indicated siRNAs were subjected to DNMT1 IP under stringent denaturing conditions. Samples were digested with trypsin, after which peptides were purified and MS was used to identify di-glycine remnants (*n*=4 independent experiments). **G.** U2OS PML-KO/YFP-PML cells were transfected with indicated siRNAs for 72 h and exposed to arsenic for 1 h. Cells were subjected to GFP IP under denaturing conditions and immunoblotted with indicated antibodies.

**Figure S5 (related to Figure 5).**
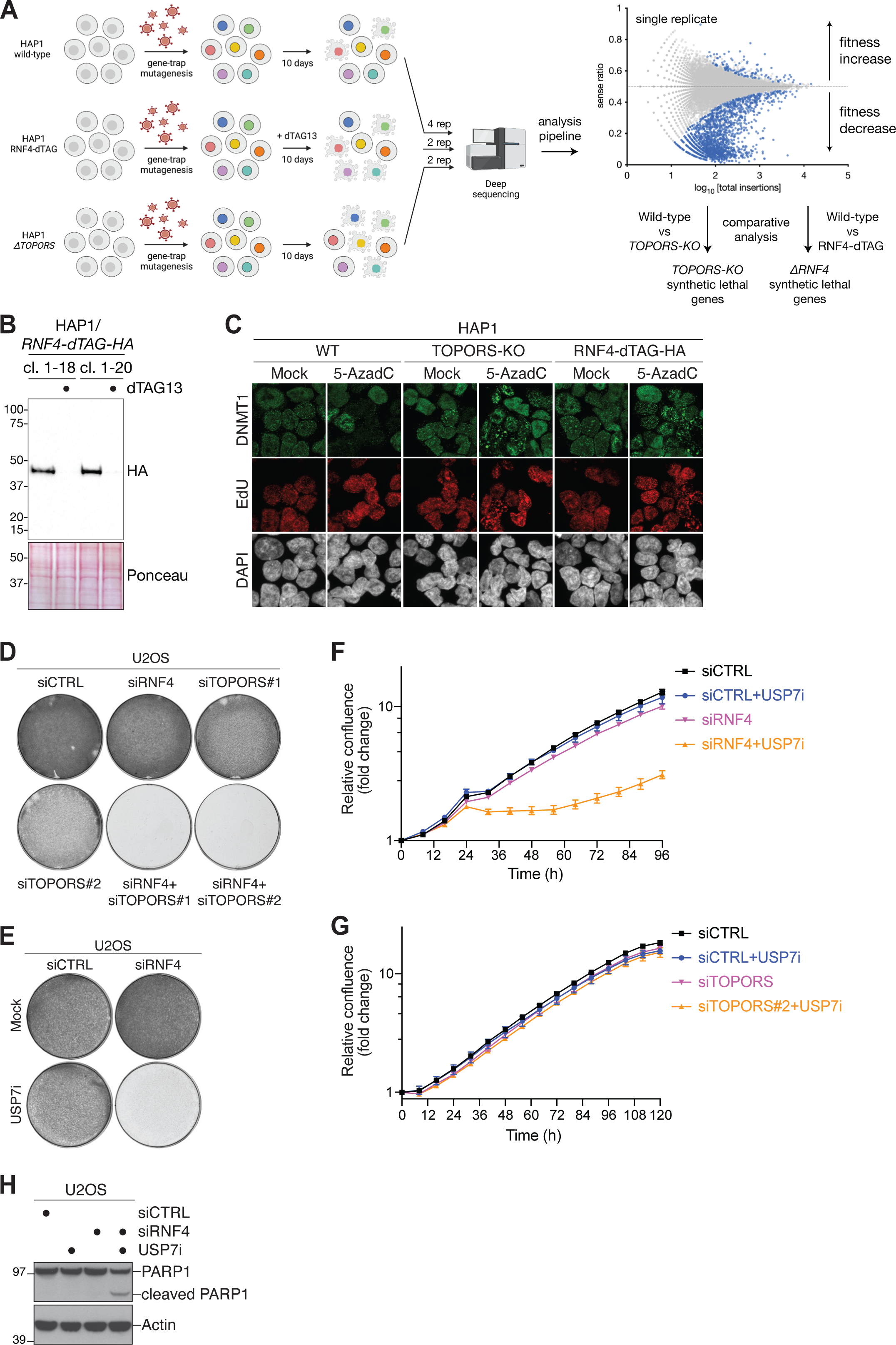
Synthetic lethality between TOPORS and RNF4. **A.** Workflow of fitness-based haploid genetic screens in Figure 5A-D. **B.** HAP1 RNF4-dTAG-HA cell line clones were treated or not with 0.25 µM dTAG13 for 24 h and processed for immunoblot analysis with HA antibody. As a loading control, the membrane was co-stained with Ponceau S. **C.** Representative images of indicated HAP1 cell lines treated with EdU in the presence or absence of 10 µM 5-AzadC for 60 min. Cells were fixed and stained using specific antibodies and reagents for DNMT1, EdU and DAPI. **D.** Equal numbers of cells was seeded in a 6-well plate, transfected with indicated siRNAs for 72 h and stained with crystal violet. **E.** As in (D), except that cells were transfected with siRNAs for 48 h and subsequently treated or not with USP7i for an additional 48 h prior to fixation. **F.** Normalized logarithmic proliferation quantification for U2OS cells transfected with non-targeting control (CTRL) or RNF4 siRNAs for 48 h and then incubated or not with USP7i, as determined by Incucyte image-based confluence analysis (mean±SD; *n*=3 technical replicates). Imaging started at 72 h after siRNA transfection. **G.** As in (F), except cells were transfected with CTRL or TOPORS siRNAs (mean±SD; *n*=3 technical replicates). **H.** Immunoblot analysis of U2OS cells transfected with indicated siRNAs for 48 h and grown in the absence of presence of USP7i for an additional 24 h.

**Figure S6 (related to Figure 5).**
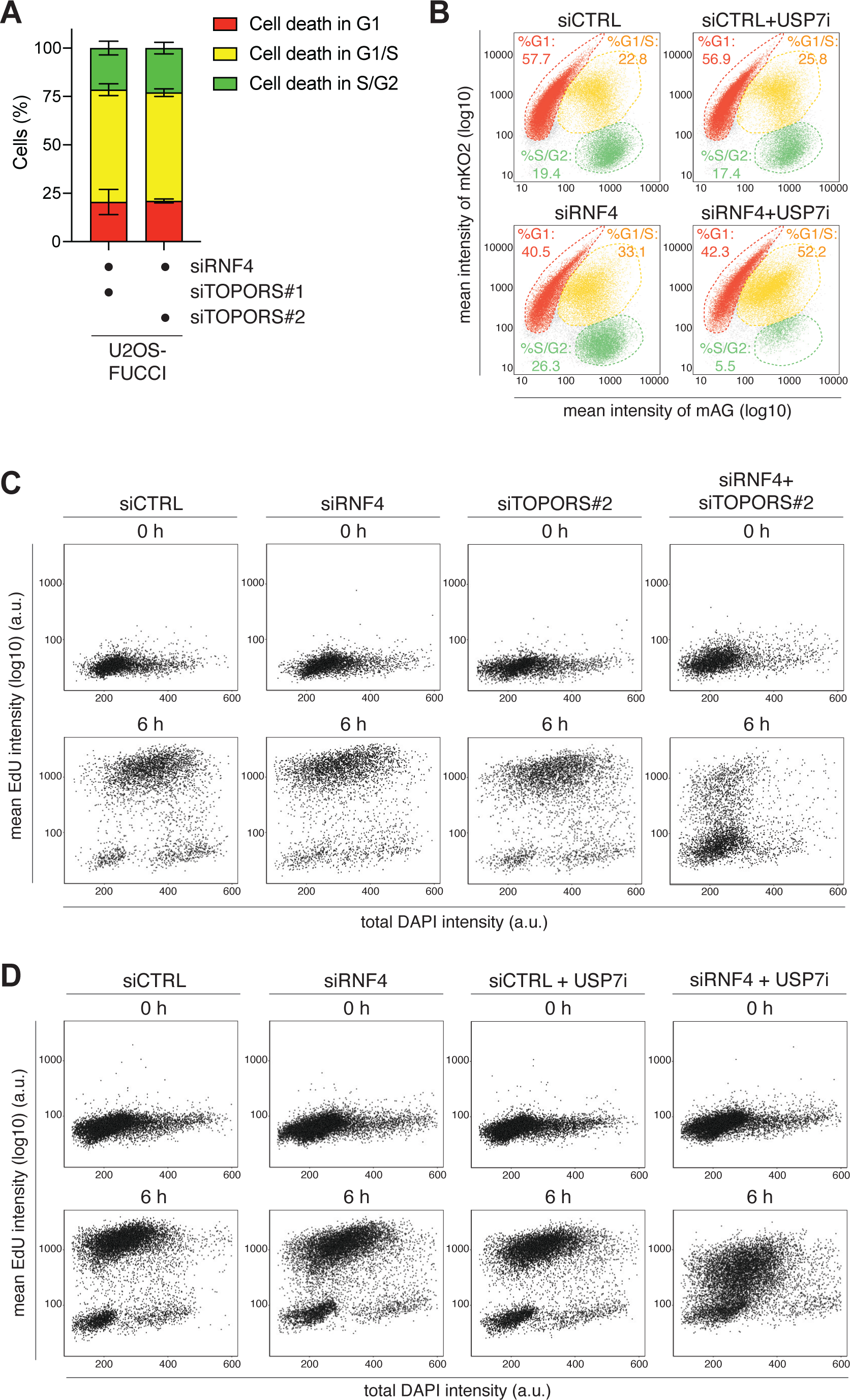
Combined loss of TOPORS and RNF4 leads to defective DNA replication and cell death in S phase. **A.** U2OS Fucci cells transfected with indicated siRNAs were subjected to live-cell imaging analysis. Expression of CDT1 (mKO2-hCdt1) and Geminin (mAG-hGem) at the point of cell death was assessed (mean±SEM; *n*=2; at least 45 cell death events were analyzed per condition per replicate). **B.** U2OS Fucci cells were transfected with indicated siRNAs for 24 h and grown in the absence or presence of USP7i for an additional 12 h. To determine cell cycle position, CDT1 (mKO2-hCdt1+) and Geminin (mAG-hGem+) intensities were analyzed by QIBC. Quantification of cells in different cell cycle phases was done using the indicated gates. Data are representative of two independent experiments. **C.** U2OS cells were transfected with indicated siRNAs and synchronized by treatment with thymidine for 18 h. DNA synthesis profiles of cells pulsed with EdU at indicated time points after release from thymidine arrest were determined by QIBC analysis of DAPI and EdU signal intensities (>3,900 cells analyzed per condition). Data are representative of two independent experiments. **D.** U2OS cells were transfected with indicated siRNAs and synchronized by treatment with thymidine for 18 h. One h prior to thymidine release, USP7i was added where indicated. DNA synthesis profiles of cells pulsed with EdU at indicated time points after release from thymidine arrest were determined by QIBC analysis of DAPI and EdU signal intensities (>10,000 cells analyzed per condition). Data are representative of two independent experiments.

**Table S1.**

**List of genes scoring as significant regulators in FACS screens for DNMT1 abundance, related to Figure 1B**

**Table S2.**

**Mass spectrometry analysis of DNMT1 IPs, related to Figure S2B, Figure 4E and Figure S4F**

**Table S3.**

**Mass spectrometry analysis of YFP-PML IPs, related to Figure 3E**

**Table S4.**

**List of significant synthetic lethal genes in HAP1 TOPORS-KO and RNF4-dTAG cells, related to Figure 5D**

## Notes

### Competing Interest Statement

The authors have declared no competing interest.

